# Sequestration of the polyunsaturated fatty acids protects the cells with oxidative phosphorylation deficiency from ferroptosis

**DOI:** 10.1101/2025.10.14.682423

**Authors:** Guillermo Puertas-Frias, María José-Saucedo-Rodríguez, Kristýna Čunátová, Lukáš Alán, Patrik Hrbáč, Michal Knězů, Soňa Koníčková, Jakub Schimmer, Monika Krakovková, Marek Vrbacký, Tomáš Čajka, Ondřej Kuda, Erika Fernández-Vizarra, Massimo Zeviani, Hana Hansíková, Tomáš Honzík, Josef Houštěk, Petr Pecina, Tomáš Mráček, Alena Pecinová

## Abstract

Impaired energy production is a hallmark of mitochondrial oxidative phosphorylation (OXPHOS) defects. However, secondary metabolic disturbances also represent an important trigger for pathologies originating from OXPHOS aberrations. We identified that cells with OXPHOS deficiencies accumulate triacylglycerols enriched in polyunsaturated fatty acids (PUFAs), which are stored in lipid droplets. Sequestration of PUFAs is a critical component of a broader stress response, which also includes downregulation of cellular desaturases and upregulation of glutathione peroxidase 4 (GPX4). Here, we demonstrate that this mechanism represents a physiologically relevant protective strategy, manifesting in the cells under hypoxia and fibroblasts derived from patients with primary mitochondrial complex IV deficiency. As proof of principle, we observed elevated PUFA-enriched triacylglycerols in the plasma of patients with Myoclonic Epilepsy with Ragged Red Fibres (MERRF). Our findings reveal a novel protective mechanism against ferroptosis, which preserves membrane integrity when mitochondrial respiration is compromised.

**Highlights:**

- OXPHOS-deficient cells trigger polyunsaturated fatty acid (PUFA) stress response that includes sequestration of PUFAs to TGs, downregulation of desaturases, and upregulation of lipid peroxide detoxification
- PUFA trafficking to TGs stored in lipid droplets protects the cells against lipid peroxidation and ferroptosis
- OXPHOS-deficient cells synthesise fatty acids de novo from glutamine-derived acetyl-CoA when reductive carboxylation is permissible
- PUFA stress response is activated in patients with mitochondrial deficiencies and during hypoxia

## INTRODUCTION

Mitochondrial oxidative phosphorylation (OXPHOS) represents a key metabolic pathway that generates cellular ATP by mitochondrial ATP synthase, which utilises the electrochemical gradient formed by the electron transport system (ETS). OXPHOS also provides intermediates for biosynthesis and maintains redox homeostasis by keeping NAD^+^/NADH balanced^1^. In humans, OXPHOS defects constitute a group of severe inborn errors of metabolism classified as mitopathies^2^. Their direct biochemical consequences depend on the specific OXPHOS component affected. However, it commonly includes decreased ATP production, accumulation of NADH, and potentially elevated levels of reactive oxygen species (reviewed in^3,4^). Accumulated NADH directly affects the activity of the tricarboxylic acid (TCA) cycle, glycolysis, malate-aspartate shuttle, and *de novo* synthesis of aspartate, which is essential for proliferation^5–7^.

To survive, cells with compromised OXPHOS function induce profound reprogramming of cellular metabolism. Increased reliance on ATP produced by substrate-level phosphorylation during glycolysis or oxidative metabolism of glutamine (TCA cycle enzyme succinyl-CoA ligase) can at least partially compensate the impaired mitochondrial ATP production^8–11^. The regeneration of NAD^+^ requires an alternative electron acceptor, such as pyruvate, which is converted to lactate^7,12^. Several groups further demonstrated that uridine is required for pyrimidine nucleotide synthesis to support proliferation when electrons cannot be transported through ETS^12,13^ to bypass dihydroorotate dehydrogenase^7,13^.

Besides upregulating glycolysis, cells with impaired ETS may redirect glutamine into the TCA cycle and convert glutamine-derived α-ketoglutarate to citrate via reductive carboxylation. It was proposed that this process is regulated by altering the α-ketoglutarate/citrate ratio^14^, but a low NAD(P)^+^/NAD(P)H ratio may facilitate the process^6^. Furthermore, in a cellular model of ATP synthase deficiency, it was demonstrated that reductive metabolism of glutamine is coupled to glycolysis via cytosolic malate dehydrogenase 1, which regenerates NAD^+15^. Another NAD^+^ regenerating mechanism to support glycolysis has been proposed by Kim and coworkers, who demonstrated that desaturase-mediated NAD^+^ recycling is an acute adaptation when mitochondrial respiration is impaired^16^.

While adaptive mechanisms for managing redox balance and generating ATP are well-documented, the downstream consequences and compensation strategies are less understood. It was demonstrated that tumour cells with mutations in subunits of complex II, a TCA cycle enzyme that is also part of the ETS, accumulate succinate, which leads to pseudohypoxia (HIF1α stabilisation under normoxic conditions)^17^. Also, in cells and tissues with compromised OXPHOS due to low levels of oxygen (hypoxia) or depletion of mitochondrial DNA, the formation of lipid droplets (LDs) was observed^18^. LDs are the lipid storage organelles that can respond to stress stimuli like nutrient and oxidative stress (e.g., due to excess lipids) by membrane remodelling^19^. Membrane phospholipids containing polyunsaturated fatty acids (PUFAs) are more prone to lipid peroxidation, and accumulation of lipid peroxides may ultimately lead to ferroptosis, a type of iron-dependent regulated cell death^20^.

PUFAs sequestration from membrane phospholipids to neutral triacylglycerols (TGs) that are stored in the core of LDs has been demonstrated in tumours^21,22^, stem cells of *D. melanogaster* exposed to hypoxia^23^ or during cell cycle arrest^24^. In contrast, lipid starvation in tumour cells resulted in PUFAs trafficking from TGs to phospholipids, promoting ferroptosis sensitivity, even though PUFA levels were decreased^25^.

Since the OXPHOS function is tightly coupled to fatty acid oxidation, we studied the remodelling of lipid metabolism as a survival strategy when OXPHOS is limited. We utilised cellular models with deficiencies in either mitochondrial complex II (CII), complex IV (CIV), or complex V (CV). We also exposed the control cells to hypoxic conditions and studied patients with mitochondrial disease. We provide evidence that cells with compromised OXPHOS activate PUFA stress response that involves the sequestration of PUFAs within neutral lipids, reduced expression of cellular desaturases, and upregulation of the antioxidant phospholipid-specific peroxidase (glutathione peroxidase 4, GPX4).

## RESULTS

### CIV-deficient cells accumulate succinate derived from glutamine

To investigate the alterations of metabolic pathways that allow cell survival in the absence of mitochondrial respiration, we employed three HEK293-derived cell lines – wild-type cells (WT), cells in which the COX6B1 subunit of CIV was knocked out (6BKO), and 6BKO cells overexpressing AOX (6BAOX)^8^, where AOX acts as the sole surrogate terminal oxidase (Figure 1A). When analysing these cells previously for the OXPHOS structural changes, we noted a secondary decrease in the levels of complex I and demonstrated that, despite abolished respiration and low NAD^+^/NADH levels, the CIV-deficient cells maintain sufficient energy charge to cover cellular ATP demands through increased glycolysis^8^.

**Figure 1:**
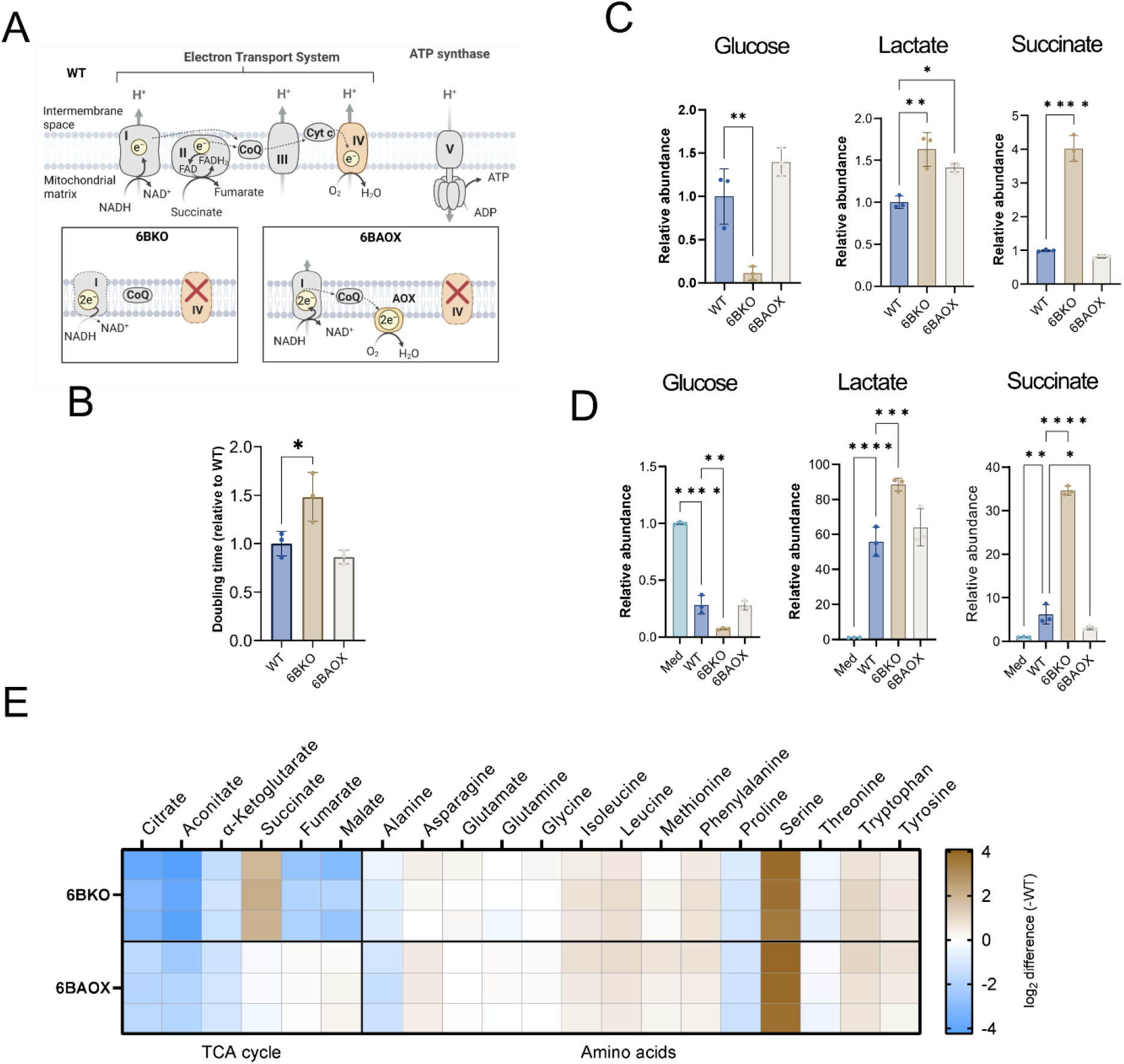
CIV deficiency promotes remodelling of the mitochondrial metabolism. (A) Scheme of the OXPHOS system in the models with knock-out of the COX6B1 subunit and the knock-out model overexpressing alternative oxidase (AOX). (B) Doubling time (relative to WT) of the cell lines measured by the CyQuant proliferation kit (n = 3). Metabolic profiling of glucose, lactate and succinate in the cells (C) and medium (D) after 24h incubation with fresh culture media. Data are expressed as fold changes to WT (C) or fresh media (D). (E) Heatmap showing fold changes in TCA cycle metabolites and amino acid levels after 24h incubation in culture media. Data represent log_2_(6BKO/WT) or log_2_(6BAOX/WT), n=3. Bar graphs represent means ± SD. One-way ANOVA (*p*-value: * <0.05; ** <0.0.1; *** <0.001; **** <0.0001) was performed.

First, we estimated the doubling time of the cell lines cultured in high-glucose medium and observed that it was significantly longer in 6BKO cells compared to WT and 6BAOX (Figure 1B). LC-MS analysis of metabolite steady-state levels revealed higher utilisation of glucose and accumulation of lactate in 6BKO cells compared to WT, both inside the cells (Figure 1C) and in the culture medium (Figure 1D). This supports our previous results, indicating that in the CIV-deficient cells, glycolysis produces ATP via substrate-level phosphorylation (SLP)^8^. Succinate was highly accumulated in 6BKO cells and media, and its level was normalised by AOX introduction in 6BAOX (Figure 1C, D, S1A). Succinate is derived from α-ketoglutarate (α-KG), which, along with upstream TCA metabolites citrate and aconitate, was significantly decreased in both 6BKO and 6BAOX cells (Figure 1E, S1A). Compared to WT, the terminal TCA metabolites fumarate and malate were lower in 6BKO and returned to control values after AOX overexpression (Figure S1A). Interestingly, 2-hydroxyglutarate, the reduced form of α-KG that was recently shown to be increased in CLPP-deficient cells^26^, was significantly elevated in 6BKO cells (Figure S1B). The levels of amino acids in 6BKO and 6BAOX were comparable to WT, except for accumulated serine (Figure 1E, S1C), which is involved in one-carbon metabolism^27^. Aspartate, essential for nucleotide synthesis^5,7^, was below the detection limit.

The data indicate that succinate is accumulated and excreted out of the cell in 6BKO cells due to the ETS dysfunction. Indeed, when electron transfer was partially restored in 6BAOX, succinate levels returned to control levels. We hypothesised that the impairment of oxidative metabolism triggers the TCA cycle to be rewired towards glutaminolysis, thereby replenishing the pools of TCA metabolites through α-ketoglutarate.

To test this, we cultured the cells for 24 hours in media containing either uniformly labelled ^13^C_6_-glucose or ^13^C_5_-glutamine, the two major contributors to the TCA cycle (Figure 2A, D), and performed fluxomic analysis. After the incubation with labelled glucose, the two carbons originating from acetyl-CoA are labelled in TCA cycle metabolites (M+2 labelled fraction) during the first turn of the TCA cycle (Figure 2A). In 6BKO cells, the M+2-enriched portion of all TCA metabolites was significantly reduced, indicating a blockage of the canonical TCA cycle. Upon expression of AOX, the levels of M+2 labelled fraction of TCA metabolites increased, although not to the levels of the WT cells (Figure 2B). A higher portion of M+3 lactate was observed in 6BKO and 6BAOX compared to WT (Figure 2C), indicating faster conversion of pyruvate to lactate and re-oxidising NADH (Figure 2A). In both 6BKO and 6BAOX cells, a significantly lower portion of pyruvate was converted to malate M+3, indicating a lesser metabolic flux via PC or ME1/2 than in the WT (Figure 2C).

**Figure 2:**
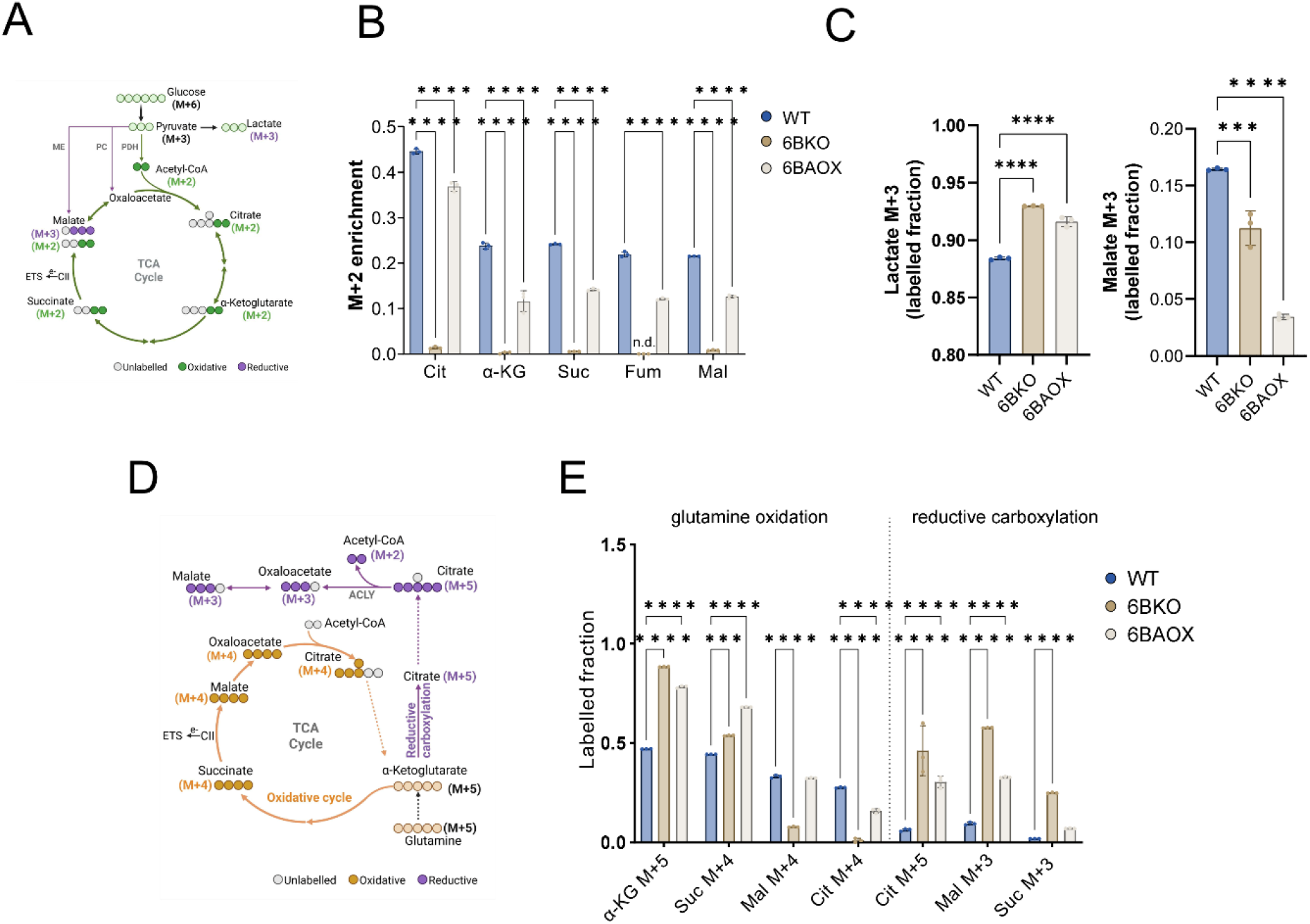
CIV-deficient cells accumulate succinate derived from glutamine. Schematic illustration of the metabolite labelling pattern from ^13^C_6_-glucose (A) or ^13^C_5_-glutamine (D). M+ indicates the number of ^13^C atoms (coloured) after 24 h incubation with the tracer. Labelled fraction (proportion of total pool) of M+2 TCA cycle metabolites (B) and M+3 lactate and malate (C) derived from ^13^C_6_-glucose. (E) Labelled fraction of the TCA cycle metabolites originating from oxidation or reductive carboxylation of ^13^C_5_-glutamine. Cit – citrate, α-KG – α-ketoglutarate, Suc – succinate, Fum – fumarate, Mal – malate, n.d. – not detected. Data are expressed as fold changes compared to WT (n=3). The bar graphs represent means ± SD (n = 3). One-way ANOVA was performed (*p*-value: * <0.05; ** <0.0.1; *** <0.001; **** <0.0001).

To track the glutamine anaplerosis of the TCA cycle (Figure 2D), we cultured the cell lines with uniformly labelled ^13^C_5_-glutamine for 24h. The fluxomic analysis revealed that approximately half of the α-KG in WT cells is derived from glutamine (α-KG M+5). At the same time, in both 6BKO and 6BAOX, the α-KG M+5 fraction is significantly increased, comprising almost the whole pool of the metabolite (Figure 2E). Succinate M+4, derived from α-KG M+5 oxidation, was increased in both 6BKO and 6BAOX models compared to WT (Figure 2E). This means that α-KG is also further metabolised in the oxidative direction (Figure 2E), allowing the substrate phosphorylation by succinyl CoA lyase^28^. In 6BKO cells, malate M+4 and citrate M+4 portions were significantly reduced, indicating that the TCA cycle is stalled at the level of succinate. AOX expression released the block in electron flux through ETS and thus also restored succinate oxidation (judged from normalised malate M+4 and citrate M+4 levels, Figure 2E). The analysis of the reductive carboxylation pathway showed a highly enriched M+5 fraction of citrate in 6BKO and, to a lesser extent, in 6BAOX, implying the elevated reductive carboxylation in both models compared to WT. The elevated levels of M+3 malate and succinate, especially in 6BKO cells, suggest that citrate is transported to the cytosol and metabolised to oxaloacetate by ATP-citrate synthase (ACLY) and subsequently converted to malate (by malate dehydrogenase). Then, malate can be transported back to the mitochondria (e.g., in exchange for citrate^29^) and enter the TCA cycle in the reverse direction to form succinate M+3 (Figure 2E).

Our data indicate that CIV-null cells accumulate succinate and 2-hydroxyglutarate, which are derived from glutamine and serve as an alternative carbon source for the TCA cycle metabolite pools. Glucose is preferentially converted to lactate and does not contribute to the TCA cycle, because it operates in the reverse (reductive carboxylation) direction.

### CIV-deficient cells accumulate triacylglycerols that can be synthesised from glutamine-derived acetyl-CoA

Conversion of citrate to oxaloacetate mediated by ACLY produces acetyl-CoA, which may become the source of carbons for *de novo* lipid synthesis. In the next step, we analysed lipid metabolism in the CIV-deficient cell line. Since mitochondrial respiration is impaired in the 6BKO model^8^, fatty acid oxidation (FAO) should not proceed. Indeed, FAO was impaired in the 6BKO model but restored by AOX overexpression (Figure 3A). Fatty acids should therefore accumulate in 6BKO. They may be elongated, desaturated, or stored directly in the form of neutral lipids.

**Figure 3:**
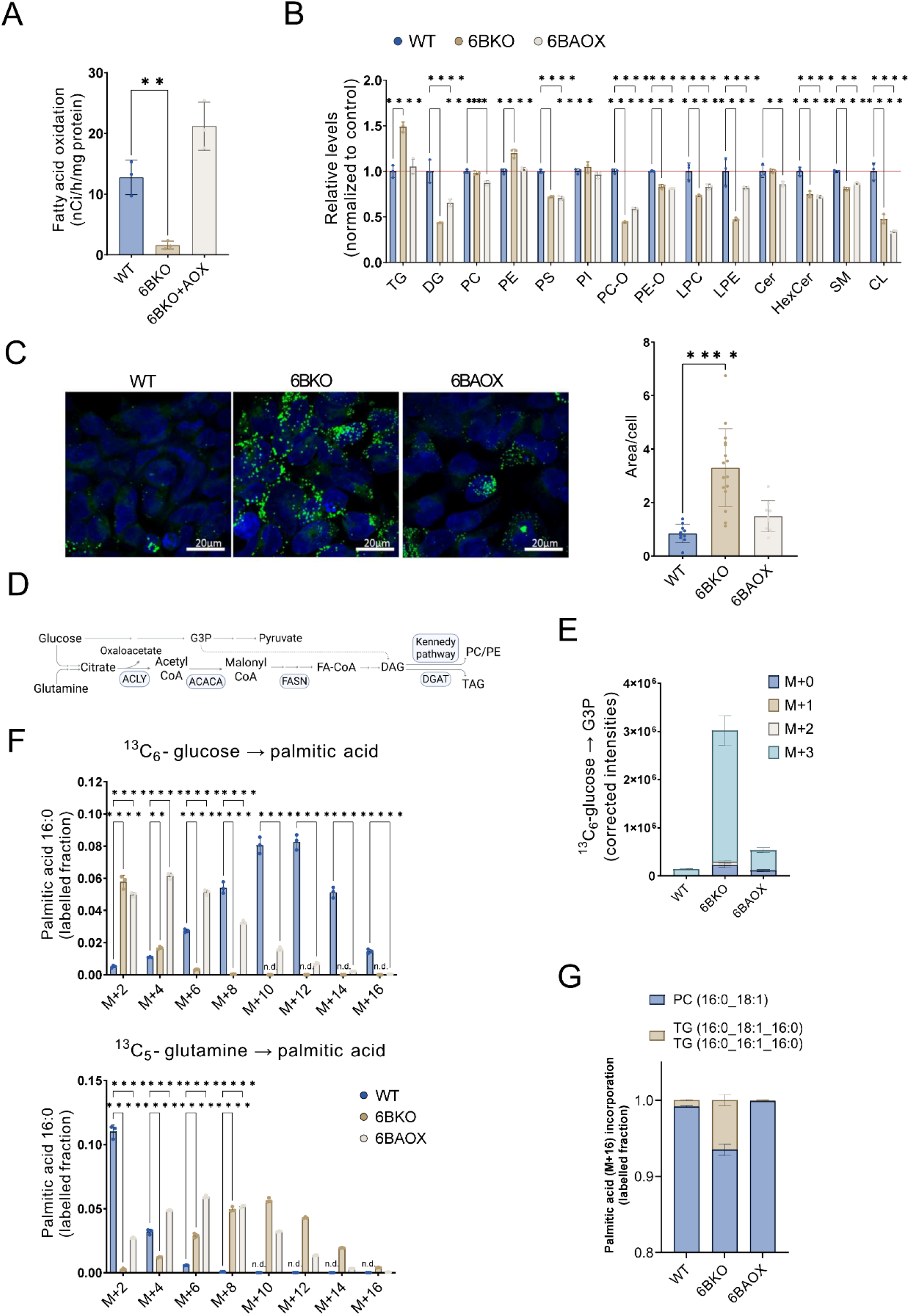
CIV-deficient cells accumulate TGs in lipid droplets and synthesise lipids *de novo* from glutamine. (A) Fatty acid oxidation of [9,10(n)-^3^H] palmitic acid (n = 3). (B) Relative levels of lipid classes measured by LC–MS (n = 3). TG-triacylglycerols, DG-diacylglycerols, PC–PC-phosphatidylcholines, PE-phosphatidylethanolamines, PS-phosphatidylserines, PI-phosphatidylinositols, PC-O-ether phosphatidylcholines, PE-O-ether phosphatidylethanolamines, LPC-lysophosphatidylcholines, LPE-lysophosphatidylethanolamines, Cer-ceramides, HexCer-hexosylceramides, SM-sphingomyelin, CL-cardiolipin. (C) Representative image and quantification of BODIPY 493/503 staining of the cell lines (n ≥ 10). (D) Schematic representation of *de novo* lipid synthesis. (E) Glycerol 3-phosphate (G3P) labelling from ^13^C_6_-glucose after 24h incubation with tracing medium (n = 3). (F) ^13^C labelling of palmitic acid after 24h incubation with ^13^C_6_-glucose or ^13^C_5_-glutamine measured by lipidomic analysis after fatty acyl chain lipolysis (n = 3). (G) Metabolic modelling of the flux of *de novo* synthesised palmitic acid into TG (1-palmitoyl-2-oleoyl-sn-glycero-3-phosphocholine; TG 16:0_18:1_18:1) or phosphatidylcholine (PC 16:0_18:1). Bar graphs represent means ± SD (n = 3). One-way ANOVA (A, C) or 2-way ANOVA (B, F) was performed (*p*-value: * <0.05; ** <0.0.1; *** <0.001; **** <0.0001).

To get a deeper insight into lipid metabolism, we performed LC-MS lipidomic analysis. The levels of triacylglycerols (TGs) were significantly increased, while diacylglycerols, TG precursors, were decreased in the 6BKO cell line, compared to WT (Figure 3B). The levels of phospholipids in 6BKO and 6BAOX were comparable to WT or slightly decreased (e.g. phosphatidylcholines (PC), or cardiolipins (CL)) except for slightly elevated phosphatidylethanolamines (PE) and reduced ether-linked phosphatidylcholines (PC-O) (Figure 3B). Microscopy analysis using Bodipy staining of neutral lipids revealed that the TGs accumulated in CIV-null cells were stored in lipid droplets (LDs)^30^ (Figure 3C). Correspondingly, glycerol 3-phosphate (G3P), which is essential for fatty acid storage in TGs (Figure 3D) was mainly derived from glucose (M+3 isotopomer) and its pool was maintained at a higher level in 6BKO than in WT cells (Figure 3E). AOX overexpression normalised all dysregulated features of TG metabolism (Figure 3B, 3C and 3E).

To uncover the origin of acetyl-CoA used for fatty acid *de novo* synthesis, we incubated the cells with labelled ^13^C_6_-glucose or ^13^C_5_-glutamine for 24h, extracted lipids, performed fatty acid hydrolysis and analysed the number of labelled carbons in fatty acyl chains. In the WT cell line, the primary source of acetyl-CoA for *de novo* palmitic (16:0) or stearic (18:0) acid synthesis was glucose, as can be seen from the higher enrichment of M+10, M+12 and M+14 palmitic and stearic acid isotopomers (Figure 3F, S2). On the contrary, acetyl-CoA in 6BKO cells was mainly derived from glutamine. 6BAOX cells then synthesise fatty acyl chains from acetyl-CoA derived from both metabolites (Figure 3F, S2). To identify whether the newly synthesised fatty acids are stored as TGs or become part of the membrane phospholipids, we analysed the labelling of lipids after 24h incubation with ^13^C_6_-glucose or ^13^C_5_-glutamine. Using metabolic modelling, we calculated the flux of *de novo* synthesised palmitic acid (16:0 M+16) to phospholipid 1-palmitoyl-2-oleoyl-sn-glycero-3-phosphocholine (PC (16:0_18:1)) or TGs 50:1 (16:0_18:1_16:0) and 48:1 (16:0_16:1_16:0) in the cell lines. Whereas labelled palmitic acid was incorporated almost exclusively into phospholipids in WT and 6BAOX, approximately 6% of *de novo* palmitic acid was deposited into TG in 6BKO (Figure 3G). Taken together, we demonstrated that in CIV-deficient cells, fatty acids are not oxidised. Still, they are synthesised *de novo*, but contrary to WT, carbons for their acyl chains originate from glutamine. Excess fatty acids, including the *de novo* synthesised ones, are stored as TGs in lipid droplets.

### Triacylglycerols accumulated in CIV-deficient cells are enriched for PUFA

Fatty acids, originating from the medium or synthesised *de novo*, can be elongated and/or desaturated. Interestingly, desaturation of fatty acids and their storage in TGs have been shown to support glycolytic NAD^+^ recycling upon acute inhibition of complex I^16^. To explore the possible role of fatty acyl chains desaturation in 6BKO, which has a low NAD^+^/NADH ratio^8^, we examined the distribution of polyunsaturated fatty acids (PUFAs) in TGs. We found that the level of TGs with a higher number of double bonds was significantly elevated in the 6BKO, as much as 9-fold in the case of specific TG species (Figure 4A). This was partially mitigated by AOX expression (Figure 4A). Based on these results and published data^16^, we hypothesised that the desaturation of fatty acyl chains is the mechanism by which 6BKO cells recycle NAD^+^ to sustain glycolysis (Figure 4B). In such a case, the inhibition of desaturases should be detrimental for the 6BKO cells. To test this, we treated the cells with inhibitors of the three major desaturases – stearoyl-CoA desaturase 1 (SCD1, inhibitor CAY10566), acyl-CoA (8-3)-desaturase (FADS1, inhibitor CP-24879) and acyl-CoA 6-desaturase (FADS2, inhibitors CP-24879 and SC-26196) and measured cellular proliferation. Surprisingly, the proliferation of the 6BKO or 6BAOX cells was not affected at 24 (Figure 4C) or 48h (Figure S3) timepoints. At the proteomic level, using label-free quantification mass spectrometry (LFQ-MS), we observed significantly decreased protein levels of the desaturases (down to a maximum 13-fold) in both 6BKO and 6BAOX cell lines (Figure 4D). This suggests that (at least in the long term) redox stress-induced PUFA formation is detrimental and counteracted through the reduction of desaturase levels. Accumulation of PUFA-rich TGs should not be considered as a mechanism of NAD^+^ recycling in 6BKO cells.

**Figure 4:**
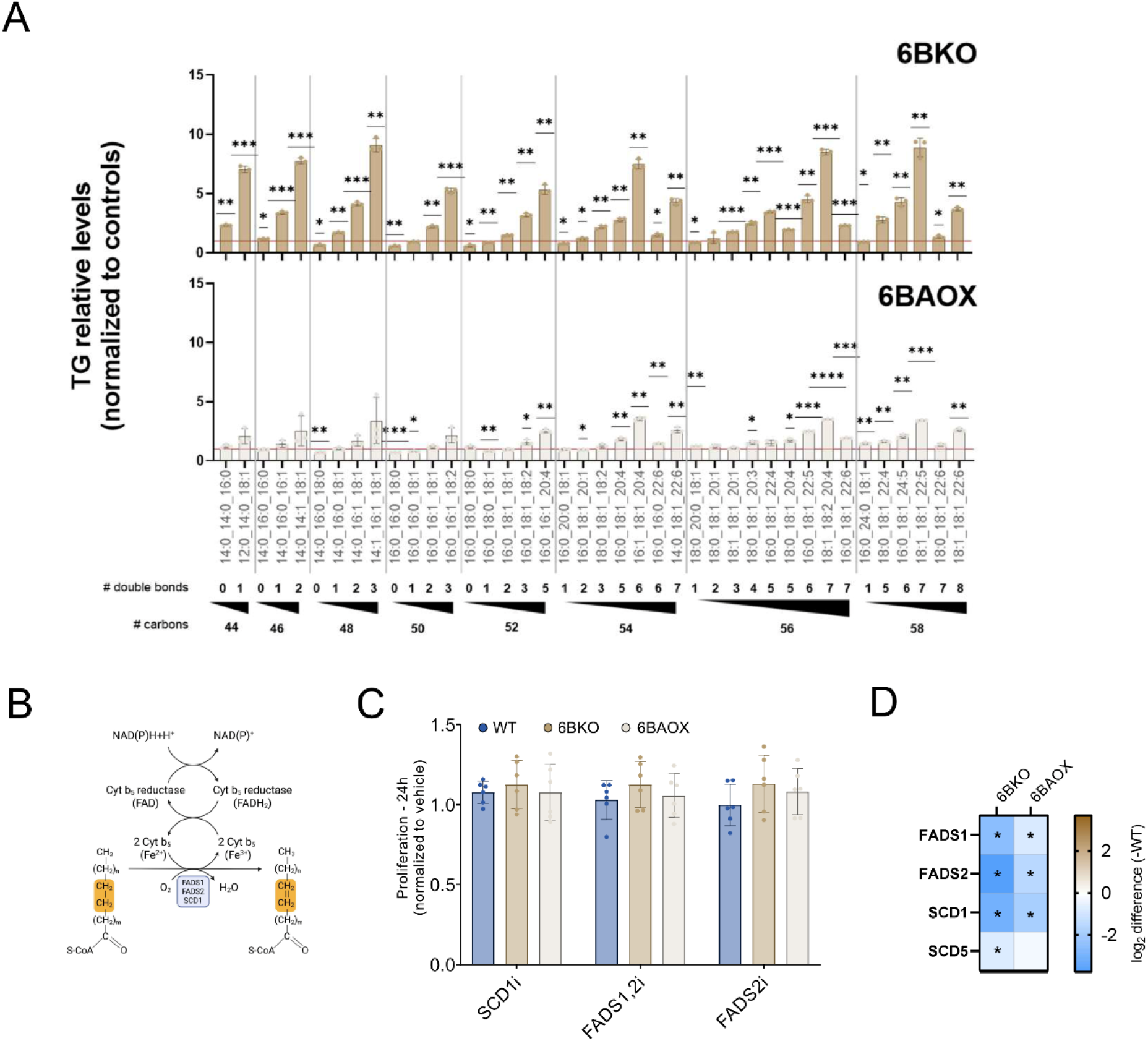
TGs are enriched for PUFAs in CIV-deficient cells. (A) Analysis of relative intracellular TGs measured by LC-MS differentiated based on the total acyl chains carbons and content of double bonds in 6BKO and 6BAOX. Data are expressed as mean ± SD, one sample t-test was performed (*p*-value: * <0.05; ** <0.0.1; *** <0.001; **** <0.0001; n = 3). (B) Scheme of fatty acyl chain desaturation. (C) Cell proliferation analysis of the cell lines incubated for 24h with inhibitors against SCD1 (CAY10566, 500nM), FADS1 and 2 (CP-24879, 30 μM) and FADS2 (SC-26196, 10 μM). Data are expressed as fold change to vehicle (n = 6). Bar graphs represent means ± SD (n = 3). 2-way ANOVA was performed (*p*-value: * <0.05; ** <0.0.1; *** <0.001; **** <0.0001). (D) LFQ-MS analysis of the cellular desaturase levels. The heatmap displays log_2_(fold change); asterisks indicate proteins that are significantly altered (*p*-value < 0.05).

### CIV-deficient cells activate PUFA-mediated stress response

Besides NADH reoxidation, higher desaturation of fatty acyl chains has been linked to a higher risk of membrane lipid peroxidation, ultimately leading to ferroptosis (Figure 5A)^31,32^. Therefore, we analysed the level of PUFAs in two major phospholipid groups – phosphatidylcholines (PC) and phosphatidylethanolamines (PE) and compared it to the level of PUFA enrichment in TGs (Figure 5B, S4A). Although some phospholipids (especially phospholipids with a higher number of double bonds) were significantly elevated by up to 40% in 6BKO compared to WT, the levels of PUFA-enriched TGs were increased as much as 4-fold in 6BKO compared to WT. Upon AOX expression, PUFAs sequestration to TGs was attenuated.

**Figure 5:**
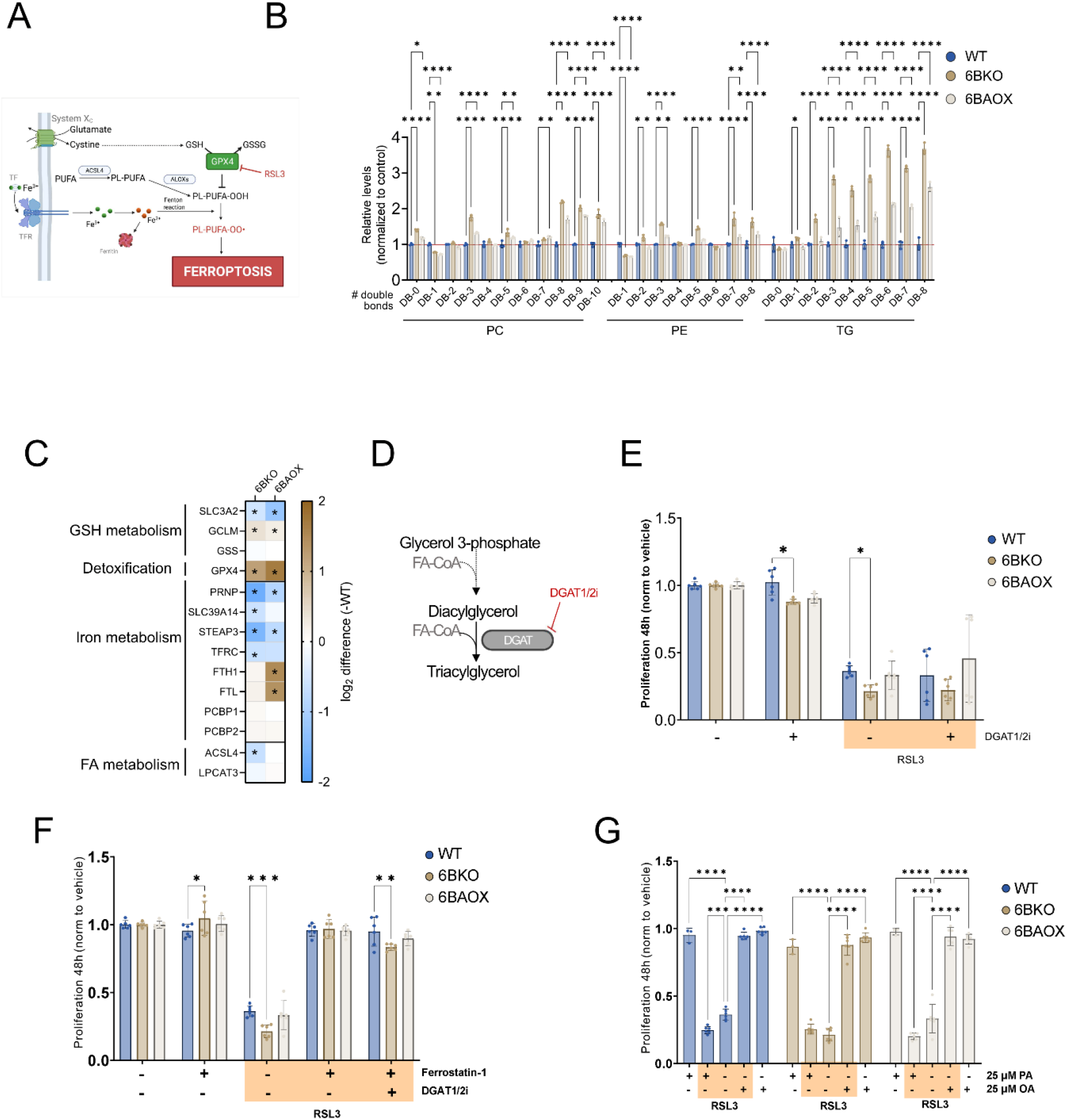
PUFA stress response is activated in CIV-deficient cells. (A) Schematic illustration of ferroptosis (according to the KEGG pathway hsa04216). (B) Relative levels of phospholipids (PC, PE) and TGs measured by LC-MS were differentiated based on double bond content. Data are expressed as fold change to WT (n = 3). (C) LFQ-MS analysis of ferroptosis-related protein levels. The heatmap displays log_2_(fold change); asterisks indicate proteins that are significantly altered (*p*-value < 0.05). (D) Schematic representation of TG formation. (E-G) Cell proliferation analysis of the cell lines incubated for 48h with an inhibitor against GPX4 (RSL3 1μM). The cells were pretreated for 24 hours with a mixture of DGAT1 (A922500, 1 μM) and DGAT2 (PF-06424439, 10 μM) inhibitors (DGAT1/2i) (E, F), the ferroptosis inhibitor ferrostatin-1 (200 nM) (F), or BSA-conjugated palmitic (PA, 25 μM) or oleic (OA, 25 μM) acid (G). Bar graphs represent means ± SD (n = 6). 2-way ANOVA was performed (*p*-value: * <0.05; ** <0.0.1; *** <0.001; **** <0.0001).

We hypothesised that the decrease of cellular desaturases and the trafficking of PUFAs into TGs in cells with deficient ETS could be a part of a PUFA-mediated stress response that maintains membrane integrity and prevents ferroptosis. A central enzyme eliminating PUFA-derived peroxyl radicals is glutathione peroxidase 4 (GPX4), which uses reduced cofactors such as glutathione (GSH) (Figure 5A, summarised in^32^). To examine whether the enzymes involved in lipid peroxidation detoxification are recruited, we performed LFQ-MS and analysed proteins involved in the ferroptotic pathway (KEGG pathway hsa04216), including GSH biosynthesis, GPX4, arachidonate/adrenic acid and iron metabolism. GPX4 levels were significantly increased in 6BKO and 6BAOX relative to WT. The changes in the abundance of proteins involved in GSH synthesis were inconsistent (Figure 5C), and GSH level did not differ among the cell lines (Figure S3B), meaning that the GSH level is not a limiting factor. The proteins involved in iron transport, such as transferrin receptor protein 1 (TFRC) or metal cation symporter SLC39A14, were decreased, while proteins involved in iron storage remained unchanged or were slightly elevated. Furthermore, the long-chain-fatty-acid--CoA ligase 4 (ACSL4), which activates fatty acids (preferentially PUFAs) that are used for membrane re-acylation by lysophospholipid acyltransferase 5 (LPCAT3), was decreased (Figure 5C), supporting our hypothesis that an excess of PUFA-enriched membrane phospholipids represents a risk for the cell.

To examine the relevance of TG formation during oxidative stress in 6BKO cells, we induced lipid peroxidation by treating the cells with the GPX4 inhibitor RSL3, a ferroptosis activator^33^. We then measured cell proliferation for 24 hours (Figure S4B-D) or 48 hours (Figure 5E-G). First, we examined proliferation when TG formation was limited by the inhibition of diacylglycerol acyltransferases (DGAT1/2i), which catalyse the terminal step of TG synthesis. We treated the cells with both the DGAT1 inhibitor A922500 and the DGAT2 inhibitor PF-06424439 (Figure 5D), which resulted in lower LD formation (Figure S4E). The inhibition of DGAT1 and 2, without inducing lipid peroxidation, resulted in a slight but significant suppression of cellular proliferation in 6BKO cells compared to WT (Figure 5E, S4B). GPX4 inhibition alone profoundly decreased proliferation in all tested cell lines; however, the 6BKO cells were significantly more sensitive than the WTs after 48 hours. Inhibition of DGAT1 and 2, in addition to the RSL3 treatment, did not further decrease proliferation in any of the models (Figure 5E). The direct cause of cell growth inhibition was the induction of ferroptosis, as the drop in cellular proliferation induced by RSL3 was prevented by the lipophilic inhibitor of lipid peroxidation, ferrostatin-1 (Figure 5F, S4C). Interestingly, the treatment with ferrostatin-1 alone slightly, yet significantly improved the proliferation of 6BKO cells compared to WT (Figure 5F). Furthermore, ferrostatin-1-mediated protection against RSL3-induced peroxidation in 6BKO cells was attenuated when DGAT inhibitors restricted TG synthesis (Figure 5F, S4C). Exogenous monounsaturated fatty acids (MUFA), unlike saturated fatty acids (SFA), have been shown to protect cells from oxidative cell death^34^. Accordingly, treatment with BSA-conjugated oleic acid (18:1) for 24 hours (Figure S4D) or 48 hours (Figure 5G) entirely blocked RSL3-induced lipid peroxidation. Treatment with palmitic acid (16:0) did not affect the phenotype in 6BKO cells but was detrimental for WT and 6BAOX cells (Figure 5G).

These results suggest that respiratory chain deficiency leads to an increase in FA desaturation, which triggers an adaptive response mitigating PUFA stress, including the sequestration of PUFAs into TGs stored in lipid droplets, the suppression of cellular desaturases, and the upregulation of lipid peroxidation protective enzymes (GPX4).

### PUFA stress response is a general phenomenon in OXPHOS-deficient cells

To investigate whether the observed PUFA stress response is exclusive to CIV-deficient cells or if it rather represents a generalised phenomenon associated with OXPHOS deficiency, we studied HEK293 cell knockouts of complex II (CII) catalytic subunit SDHA (SDHAKO) and complex V (CV) catalytic subunit ATP5F1B (BETAKO). Both knockout cell lines lack the assembled CII or CV, respectively, and exhibit a secondary CI deficiency^8^. Out of the OXPHOS deficiency models, SDHAKO had the highest cellular oxygen consumption rate (OCR), but all of them (SDHAKO, BETAKO, 6BKO) respired significantly less than WT (Figure S5A). The glycolytic activity, estimated from extracellular acidification rate (ECAR), was elevated compared to WT in all tested cell lines, indicating increased glycolysis and lactate production (Figure S5A). LC-MS analysis of TCA cycle metabolites in SDHAKO revealed an accumulation of succinate and a significant decrease in all the other TCA metabolites, whereas in BETAKO cells, only citrate and aconitate levels were significantly decreased (Figure 6A, S5B). The levels of amino acids in SDHAKO and BETAKO cells mirrored those in 6BKO cells, with a prominent increase in serine levels.

**Figure 6:**
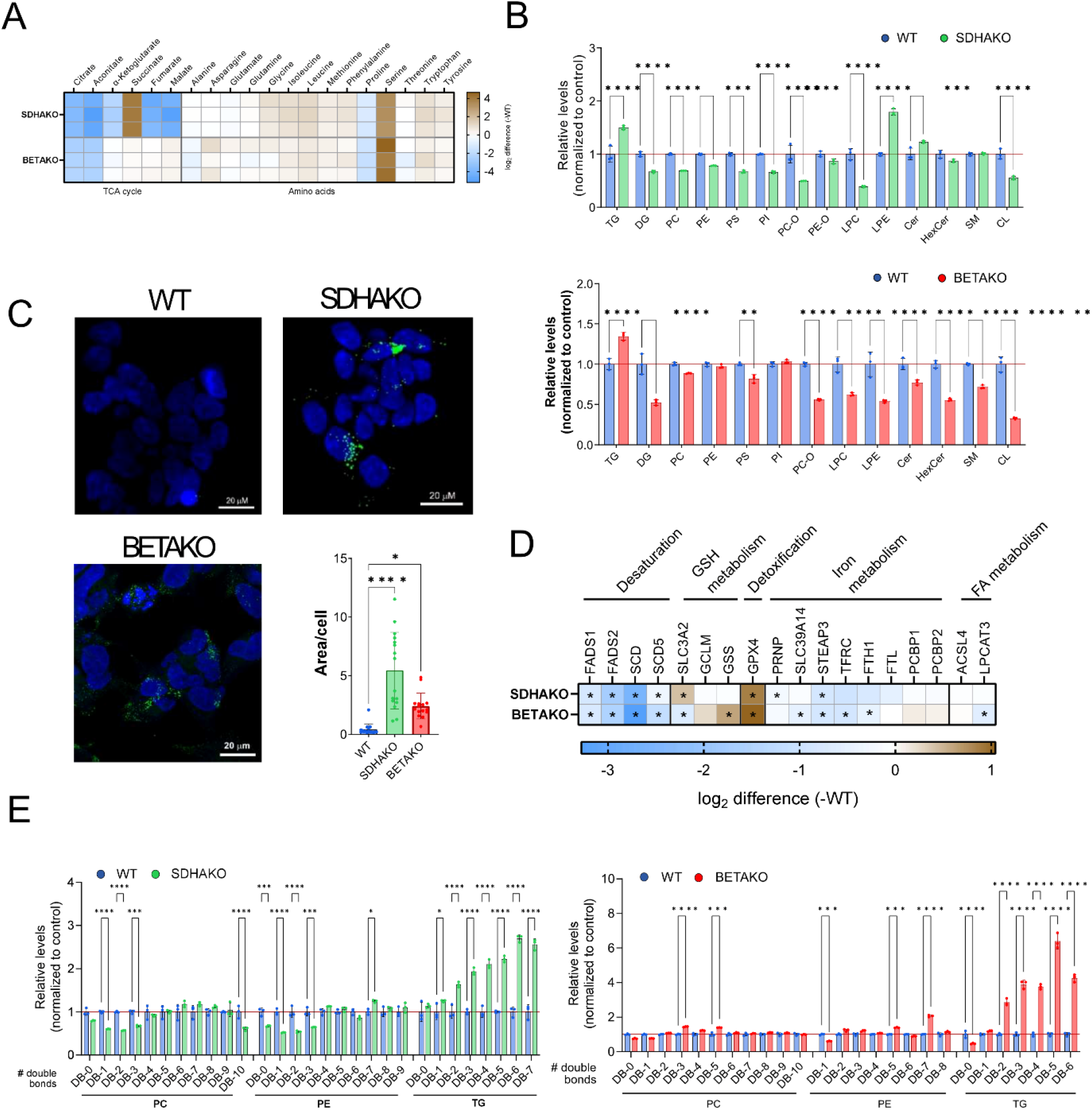
OXPHOS dysfunction leads to PUFA stress response. (A) Heatmap showing fold changes in TCA cycle metabolites and amino acid levels after 24h incubation in culture media. Data represent log_2_(fold change) of SDHAKO or BETAKO compared to WT (n = 3). (B) Relative levels of lipid classes measured by LC–MS (labelled as in Figure 3B). (C) Representative image and quantification of BODIPY 493/503 staining of the WT, SDHAKO and BETAKO cell lines (n ≥ 15). (D) LFQ-MS of ferroptosis-related protein levels. The heatmap displays log_2_(fold change) of SDHAKO or BETAKO compared to WT (n = 3). Asterisks denote proteins that have changed significantly (*p*-value < 0.05). (E) Relative levels of phospholipids (PC, PE) and TG measured by LC-MS differentiated based on double bond content (n = 3). Bar graphs represent means ± SD. One-way (C) or 2-way ANOVA (B, D, E) was performed (*p*-value: * <0.05; ** <0.0.1; *** <0.001; **** <0.0001).

SDHAKO and BETAKO cells recapitulated the features observed in 6BKO: TGs were accumulated (Figure 6B), stored in lipid droplets (Figure 6C), levels of cellular desaturases were suppressed, and GPX4 was elevated (Figure 6D). Furthermore, PUFAs were enriched in TGs to a greater extent than in phospholipids, suggesting they were sequestered into TGs (Figure 6E, S5C, D). ^13^C_6_-glucose and ^13^C_5_-glutamine tracing analysis showed that the flux of glucose-derived metabolites through the TCA cycle was decreased in BETAKO cells and was only negligible in SDHAKO (Figure S6A). Compared to WT cells, α-KG M+5 and succinate M+4, derived from glutamine oxidation, were accumulated in both models, implying an increased flux in the oxidative direction of the TCA cycle. The labelled fraction of glutamine-derived malate M+4 was decreased in BETAKO and barely detectable in SDHAKO cells (Figure S6B). BETAKO cells resembled 6BKO in their handling of glutamine. We observed increased metabolic flux of glutamine through reductive carboxylation (elevated citrate M+5 and malate M+3, Figure S6B) and preferential use of glutamine-derived acetyl-CoA to synthesise palmitic acid *de novo* (Figure S6D). This contrasted with SDHAKO cells, where we observed only a slight increase in the rate of reductive carboxylation (assessed by labelling of malate M+3, Figure S6B) and subsequently a moderate increase in the contribution of glutamine-derived acetyl-CoA to the *de novo* synthesis of palmitic acid (Figure S6C). To investigate how reductive glutamine metabolism is regulated in our models, we assessed the NAD^+^/NADH ratio and the α-KG/citrate ratio, as these are the two proposed regulatory mechanisms^6,14^. When assessing the NAD+/NADH ratio, we found a significant decrease in 6BKO, which was normalised upon AOX overexpression. In BETAKO and SDHAKO cells, this ratio was comparable to that in WT cells (Figure S6E and our previous data^8^). On the other hand, the α-KG/citrate ratio measured by LC-MS was the most increased in 6BKO and BETAKO cells (more than four-fold) and only to a lesser extent in SDHAKO cells (Figure S6F). These data indicate that a high α-KG/citrate ratio is a more important driver of reductive glutamine utilisation in OXPHOS-deficient cells.

In conclusion, the PUFA stress response is also activated in CII- or CV-deficient cells, and the sequestration of PUFAs into neutral lipids stored in LDs represents a common feature among OXPHOS-deficient cells. While de novo-synthesised FAs contribute to the pool of accumulated TGs, they are not an exclusive prerequisite of PUFA stress activation. PUFA stress response thus involves FAs derived from multiple sources.

### Hypoxia triggers the PUFA stress response

The inefficiency of oxidative phosphorylation and the formation of lipid droplets have been previously associated with hypoxia^18,35^. To investigate whether the PUFA stress response is activated during hypoxia, we cultured WT cells in a 1% O_2_ atmosphere for one week. TGs together with lysophosphatidylcholines (LPC) levels (Figure 7A), as well as the lipid droplet area (Figure 7B), were increased in hypoxia compared to normoxia (21% O_2_). Under hypoxia, PUFAs were mainly enriched in the accumulated TGs and, to a lesser extent, in phospholipids (PC and PE) (Figure 7C). LFQ-MS analysis revealed reduced protein levels of desaturases (FADS1 and 2, SCD) and elevated GPX4 (Figure 7D). Therefore, the PUFA stress response is an adaptive mechanism that is triggered even when OXPHOS function is impaired due to limited oxygen levels.

**Figure 7:**
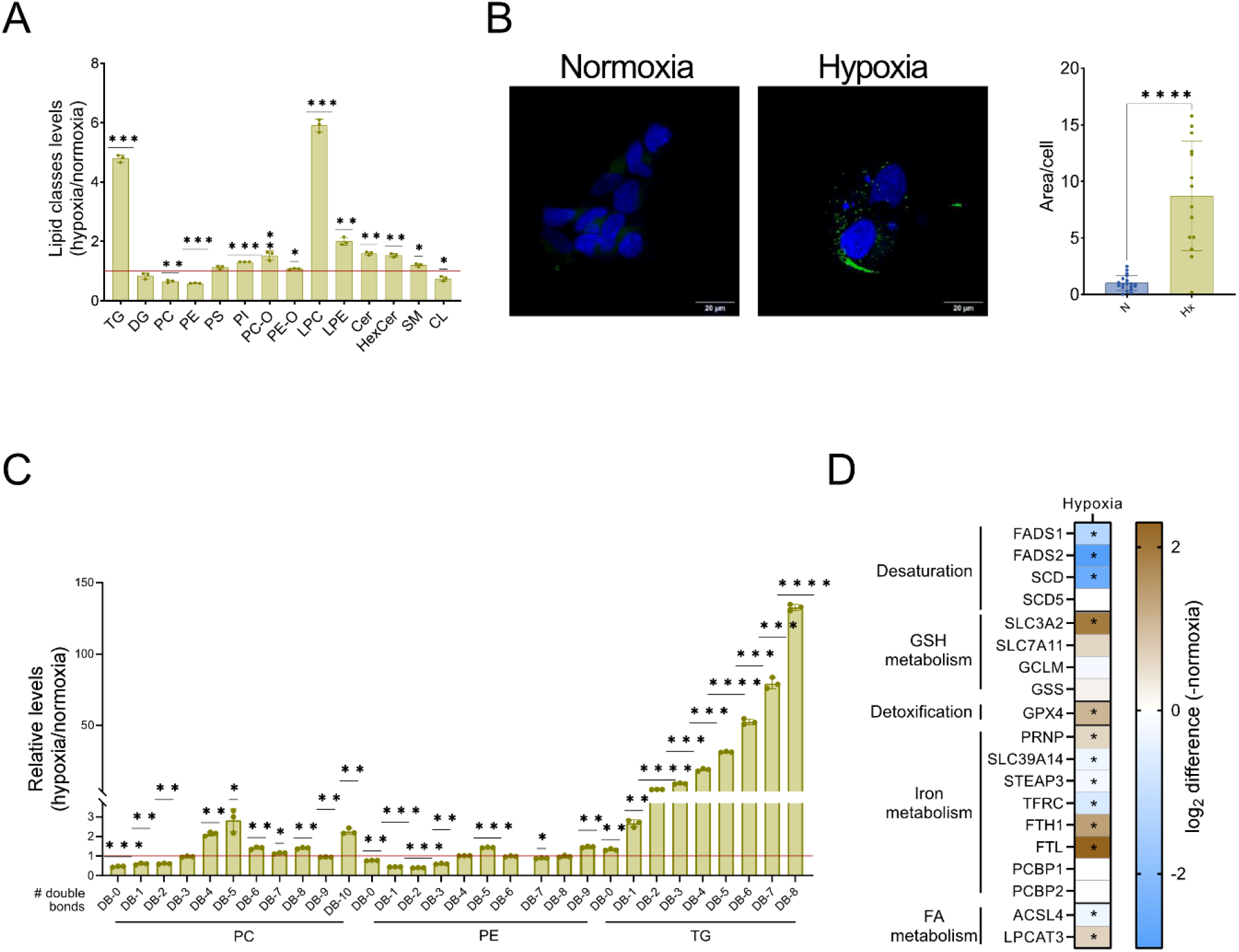
Hypoxia treatment induces PUFA stress response in WT cells. (A) Relative levels of lipid classes measured by LC–MS (labels as in Figure 3B). (B) Representative image and quantification of BODIPY 493/503 staining of the WT in normoxia or hypoxia (n ≥ 15). (C) Relative levels of phospholipids (PC, PE) and TG measured by LC-MS were differentiated based on double bond content (n = 3). (D) LFQ-MS analysis of ferroptosis-related protein levels. The heatmap displays log_2_(fold change) of the values in hypoxia compared to normoxia (n = 3). Asterisks denote proteins that have changed significantly (*p*-value < 0.05). Bar graphs represent means ± SD. One-way (B) or 2-way ANOVA (A, C) was performed (*p*-value: * <0.05; ** <0.0.1; *** <0.001; **** <0.0001).

### PUFA stress response is an adaptive mechanism in patients with mitochondrial deficiency

Next, we investigated whether PUFA stress response activation is relevant to human pathology in cases of mitochondrial deficiencies. First, we analysed the immortalised human skin fibroblasts derived from two patients suffering from leukodystrophy and carrying a missense variant in the CIV structural subunit COX6B1 (p.R20H)^36^. The biochemical activity of cytochrome *c* oxidase was significantly decreased in the patient fibroblasts compared to the control (Figure 8A)^36^. Lipid class evaluation showed increased TG levels in the patient cells (Figure 8B), which also accumulated lipid droplets (Figure 8C). Similar to the OXPHOS dysfunction models investigated above, patient fibroblasts displayed reduced protein levels of fatty acid desaturases (SCD, FADS1, and FADS2) and elevated GPX4 (Figure 8D). PUFA sequestration analysis phenocopied the results from the HEK293 KO models, displaying highly elevated PUFAs within TGs and, to a lesser extent, in phospholipids (PC and PE, Figure 8E).

**Figure 8:**
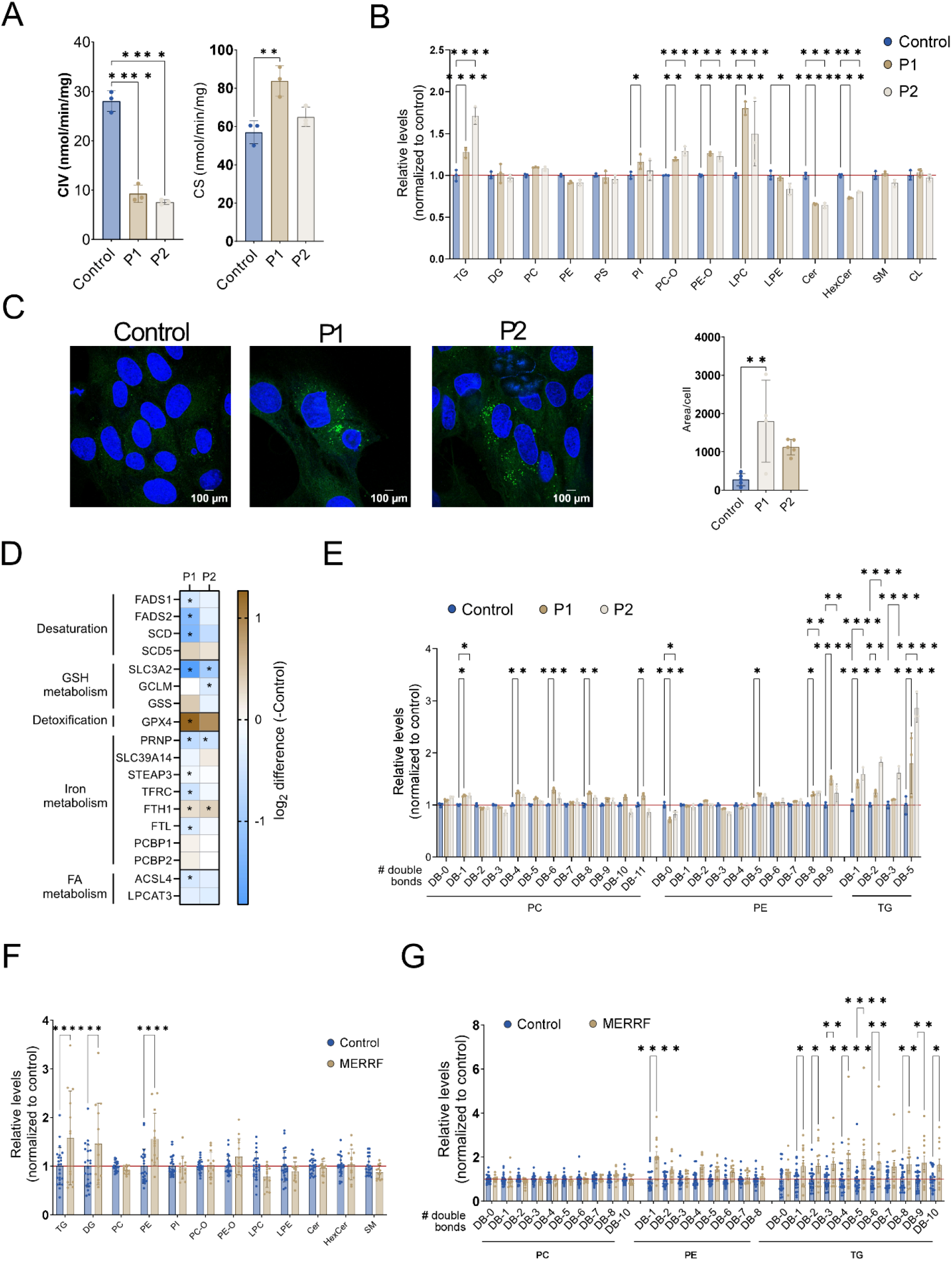
PUFA stress response is activated in patients with mitochondrial deficiency. (A) Cytochrome *c* oxidase (CIV) and citrate synthase (CS) enzyme activities measured in immortalised fibroblasts (control and derived from patients with the mutation in the *COX6B1* gene – P1, P2) were assessed by spectrophotometer (n = 3). (B) Relative levels of lipid classes measured by LC–MS (labelled as in Figure 3B), n = 3. (C) Representative image and quantification of BODIPY 493/503 staining of the control and the patients’ cells (n ≥ 4). (D) LFQ-MS analysis of ferroptosis-related protein levels. The heatmap displays log_2_(fold change) of the values in P1 or P2 compared to control (n = 3); asterisks denote proteins that have changed significantly (*p*-value < 0.05). (E) Relative PC, PE, and TG levels differentiated based on the double bond content measured by LC–MS (labelled as in Figure 3B) in control and patients’ fibroblasts (n = 3). Relative levels of lipid classes (F) and PC, PE and TG (G) differentiated based on the content of double bonds measured by LC–MS (labelled as in Figure 3B) in plasma of controls (n = 23) and MERRF patients (n = 13). Bar graphs represent fold changes of patients and control values. Bar graphs represent means ± SD. One-way (A, C) or 2-way (B, E – G) ANOVA was performed (*p*-value: * <0.05; ** <0.0.1; *** <0.001; **** <0.0001).

To elucidate whether our observations can also be valid *in vivo*, we compared plasma samples from control individuals and patients with myoclonic epilepsy with ragged red fibres (MERRF) syndrome. MERRF is a mitochondrial disease that typically affects the muscles and nervous system^37^. In accordance with a previous study^38^, the analysis of lipid classes demonstrated an increase in TGs and DGs (Figure 8F). This indicates an overload of fatty acids and/or an upregulation of lipid synthesis pathways. The specific rise of PE in the blood may indicate altered turnover or degradation of PE, for example, during mitophagy, as the PE content in mitochondrial membranes exceeds 30 %^39^. The detailed analysis of phospholipids and TGs showed an accumulation of PUFA-enriched circulating TGs in MERRF patients relative to controls (Figure 8G). Taken together, the data from COX6B1-deficient patient-derived fibroblasts and plasma from individuals with MERRF syndrome suggest that the PUFA stress response is a mechanism relevant to mitochondrial disease.

## DISCUSSION

Mitochondrial OXPHOS defects impose a profound metabolic stress, forcing cells to reprogram bioenergetic and biosynthetic pathways to survive. While compensatory mechanisms for ATP production and NAD^+^ regeneration are becoming increasingly well-characterised^7–12^, the downstream metabolic consequences remain less clear. Here, we identified a polyunsaturated fatty acid (PUFA) stress response as a unifying adaptive mechanism in OXPHOS-deficient cells, characterised by sequestration of PUFAs into triacylglycerols (TGs) stored in lipid droplets (LD), suppression of desaturase expression, and upregulation of glutathione peroxidase 4 (GPX4). This adaptive response was observed across distinct genetic models of complex II, IV, and V deficiencies, during hypoxia, and in primary fibroblasts from mitochondrial disease patients. In addition, we identified PUFA-enriched TGs in the plasma of patients with MERRF syndrome.

It has been shown that PUFAs are highly susceptible to oxidative damage^40,41^ and their accumulation in membrane phospholipids can increase sensitivity to ferroptosis, even when overall PUFA levels are low due to lipid limitation^25^. Our findings demonstrate that cells with compromised OXPHOS trigger a multi-pronged defence strategy to reduce PUFA content in the membrane phospholipids. First, the upregulation of GPX4, the enzyme that directly reduces phospholipid hydroperoxides, has been shown to confer resistance to ferroptosis. In contrast, its knockdown increased sensitivity to various ferroptosis inducers^42^. In cancer, a high level of GPX4 has been associated with drug resistance^43^ and correlated with resistance to ferroptosis^44,45^. In mice, inducible GPX4 upregulation protected retinal cells from oxidative damage^46^. The importance of GPX4 has also been illustrated during adipocyte differentiation^47^, spermatogenesis^48^, ischemia/reperfusion injury^49,50^ or in metabolic-associated fatty liver disease^51^. Notably, GPX4 protein level was increased in a heart-specific mouse model of complex IV deficiency by a transcriptional activation mechanism mediated by Nrf2 and Atf4^52^. Interestingly, the Atf4 protein level is increased in 6BKO and BETAKO cells^8^ suggesting a possible mechanism of GPX4 regulation.

The next level of the defence mechanism is lowering the rate of fatty acid desaturation by decreasing the expression of fatty acid desaturases. The regulation of the expression of the three primary desaturases (SCD1, FADS1 and 2) is complex, involving changes in various transcription factors (such as SREBP-1), hormonal signalling and dietary carbohydrates or fatty acids (summarised in^53^). It has been published that PUFAs reduced FADS1 and FADS2 gene expression in differentiated 3T3-L1 cells^54^, which could represent a possible negative-feedback mechanism to decrease their activities in OXPHOS-compromised cells.

The mode of regulation of desaturases was previously explored under the conditions of acute OXPHOS dysfunction. It was published that complex I inhibition (24h treatment with rotenone) in cultured mouse renal epithelial cells resulted in the accumulation of highly unsaturated fatty acids (HUFAs) stored mainly in TGs. It was shown that the activity (not the amount) of the FADS1 and FADS2 desaturases is increased and that HUFA formation helps to recycle glycolytic NAD^+16^. In the mouse model, rotenone treatment for 10 hours resulted in the accumulation of HUFA-enriched TGs in the kidney and liver, while the protein level of Fads1 was unaffected; however, mRNA levels of *Fads1* and *Fads2* were decreased^16^.

In our model of sustained OXPHOS deficiency, the desaturases are among the most downregulated proteins (e.g., FADS2 is decreased 3–13-fold in KO models), indicating the inhibition of PUFA formation. At the same time, under the conditions of compromised fatty acid oxidation, cells activate lipid remodelling to maintain a balanced phospholipid composition in the membranes. This suggests that, as a long-term adaptation, cells prioritise survival by suppressing the production of PUFA by desaturases, even at the cost of compromised NAD+ recycling through fatty acid desaturation. Other enzymes involved in NAD^+^ recycling, such as lactate dehydrogenase^7,12^ or malate dehydrogenase 1^15^, have to maintain a survival-permissive NAD^+^/NADH redox ratio.

We have also observed a PUFA stress response in SDHAKO cells, where blockade of TCA cycle flux might be expected to shift the NAD^+^/NADH ratio to an even more oxidised state. Indeed, this has been observed previously, when complex II inhibition led to an increase in NAD^+^/NADH, which was normalised when complex I was also inhibited^55^. Furthermore, the authors observed an adaptive decrease in complex I in SDHB-null cells, which enabled a higher proliferation rate. This can also be observed in our SDHAKO cells, where downregulated complex I coincides with a normalised NAD^+^/NADH ratio^8^. OXPHOS system disorders in general are associated with redox imbalance, manifesting either as overreduction of NAD^+^/NADH or an increase in mitochondrial ROS production (observed for CII^56^ or CV^8,57,58^ defects) and ultimately results in irreversible oxidation of macromolecules, such as lipid peroxidation^59^. Abnormal flux through the respiratory chain is also likely the driver behind PUFA formation in the mitochondrial membrane, which has to be counteracted by the PUFA stress response reported in this paper. Interestingly, Sonlicromanol, which recently secured Orphan Drug Designation for mitochondrial OXPHOS defects, acts as a cellular redox status modulator and attenuates lipid peroxidation^60,61^. It is therefore likely that PUFA stress response represents a molecular pathway that targets patients’ cells and tissues.

The final level of the cellular response involves the sequestration of PUFAs into TGs stored in LDs. Since phospholipids containing PUFAs are the most susceptible to oxidation^62^, this process protects cellular membrane integrity. LD formation has been associated with dysregulated lipid metabolism, and LD accumulation has been well documented in a broad range of malignant cancers, under hypoxia-induced OXPHOS deficiency, in cells lacking mitochondrial DNA, and in other conditions (summarised in^19^). The role of lipid droplets as antioxidant and pro-survival organelles that retain PUFAs in TGs and thus protect the cells from lipotoxic stress has been demonstrated in breast cancer^22^, in an acidic tumour environment^21^, during neuronal development in *D. melanogaster*^*23*^ and cell cycle arrest^24^. Here, we have demonstrated that the sequestration of PUFAs to TGs stored in lipid droplets is a common feature in cells with sustained OXPHOS insufficiency and that the PUFAs trafficking is a part of a complex PUFA stress response. Recently, a reverse process of PUFA trafficking from TGs to membrane phospholipids has been observed in cancer cells with depleted extracellular lipids, despite reduced levels of cellular PUFAs^25^. The lipid-starved cells liberated PUFAs from TGs, converted them to HUFA and used them to synthesise phospholipids, thus increasing cancer cell sensitivity to ferroptosis. These findings, along with our data, suggest that dynamic communication between LDs and the membrane occurs when lipid metabolism is dysregulated, enabling cells to regulate membrane composition actively, thus preserving membrane integrity.

Proper OXPHOS function is essential for many vital cellular processes, including mitochondrial ATP production, redox status regulation, and synthesis of macromolecules^63^. Previous studies have demonstrated that substrate-level phosphorylation covers cellular ATP demands in cells with CII, CIII, or CV deficiency^6,9,15,64,65^. In agreement, our data show that cellular ATP requirements are covered by increased glycolysis, as evidenced by increased glucose consumption and lactate production in 6BKO, SDHAKO, and BETAKO cells. Furthermore, mitochondrial respiration has been shown to be essential for cellular proliferation as it enables aspartate synthesis^5,7^. In our KO models, the aspartate steady-state level could not be assessed because it was under the detection limit. However, our fluxomic data indicate that oxaloacetate, a precursor of aspartate, is produced via malate dehydrogenase or ATP-citrate synthase. This suggests that any aspartate produced via these pathways is rapidly consumed, allowing the proliferation of KO cells. Additionally, our analyses of cellular metabolism revealed that glucose oxidation is impaired and succinate is accumulated in 6BKO and SDHAKO cells, similar to what has been described in other types of OXPHOS dysfunction^6,15,64–67^. Moreover, in all tested models, the serine level was significantly increased. Recently, it has been reported that de novo serine biosynthesis may protect against mitochondrial failure in skeletal muscle^68^. Conversely, serine catabolism requires NAD^+ 69^; therefore, it might be hampered in our cell models. Another process linked to NAD^+^ might be glutamine reductive carboxylation. Its increase has been frequently observed in cells with mitochondrial dysfunction^6,14,15,55,65,67^ or in hypoxia^14,66,70^. It was proposed that this process is regulated by the NAD^+^/NADH ratio^15,67^ and/or the α-ketoglutarate/citrate ratio^14^. In all our OXPHOS-deficient models, reductive glutamine metabolism was running, but the flux through the pathway differed. While approximately 40-60% of citrate and malate was derived from glutamine in 6BKO and BETAKO, only 20% of glutamine-derived malate (labelled citrate was under the detection limit) was observed in SDHAKO cells. Since the NAD^+^/NADH ratio was reduced only in 6BKO cells, we can conclude that reductive carboxylation in OXPHOS-deficient HEK cells is rather regulated by the α-ketoglutarate/citrate ratio, in accordance to what was proposed in^14^.

We also provide evidence that glutamine-derived acetyl-CoA is used for *de novo* lipogenesis, especially in 6BKO and BETAKO cells, and that in OXPHOS-deficient cells, the newly synthesised fatty acids are incorporated into TGs to a higher extent. Interestingly, increased glutamine reductive carboxylation and incorporation of glutamine-derived carbons into fatty acids have also been demonstrated in A549 cells exposed to hypoxia^66^.

Together, these observations suggest a coordinated reprogramming of lipid metabolism to decrease lipid peroxidation, thus preserving membrane integrity and protecting cells against ferroptosis. In addition, we have demonstrated that the PUFA stress response is not limited only to severe OXHOS deficiency KO cell models. We have found evidence that it is also relevant in mitochondrial CIV-deficiency patient-derived fibroblasts and, based on the analysis of plasma lipidome, most likely also in patients with MERRF syndrome. Elevated PUFA-rich TGs in circulation may reflect systemic adaptation to mitochondrial dysfunction and serve as a biomarker for diseases characterised by limited OXPHOS function, as supported by our current data. Previous studies in mitochondrial disease patient cohorts have reported lipid dyshomeostasis in plasma^71^. Our work directly links these changes to protection from ferroptosis; however, larger cohorts are required to validate PUFA-rich TGs as a biomarker. Our data also suggests that therapies using DGAT inhibitors, which aim to reduce steatosis, could be harmful in the context of mitochondrial disease. Conversely, a diet enriched with MUFA, such as oleic acid, could reduce the risk of ferroptosis^34^ but increases the risk of liver disease^72^.

### Limitations of the study

The experimental work was performed primarily in *in vitro* cell cultures, accompanied by metabolomic analysis of plasma samples from patients with mitochondrial deficiencies. The next step would be to validate the findings in animal models of OXPHOS limitation (e.g. a mouse model of OXPHOS defect or mice exposed to hypoxia). Investigating the upstream signalling pathways that initiate lipid remodelling in OXPHOS-compromised cells would also be an important future direction.

## Acknowledgements

This work was supported by the Czech Science Foundation (GACR 21-18993S; GP, KC, LA, MV and AP), the National Institute for Research of Metabolic and Cardiovascular Diseases (Programme EXCELES, ID Project No. LX22NPO5104) – Funded by the European Union-Next Generation EU (JH and TM), the Czech Health Research Council (NU22-01-00499; HH, TH and PP) and the Grant Agency of Charles University (GA UK 283423/2023, GP and MJS). The authors would like to acknowledge the Laboratory of Metabolomics at the Institute of Physiology of the Czech Academy of Sciences (TC) and the Proteomics Service Laboratory at the Institute of Physiology (supported by RVO, ID 67985823) and the Institute of Molecular Genetics (supported by RVO, ID 68378050) of the Czech Academy of Sciences (MV). Imaging was supported by IPHYS BIF – MEYS CR (Large RI Project LM2023050 Czech-BioImaging) and ERDF (Project No. CZ.02.1.01/0.0/0.0/18_046/0016045). We sincerely thank Barbora Opletalova and Marketa Hlavackova for their expert technical support, which was essential for the successful execution of the experiments within the hypoxic chamber.

## Author contributions

Conceptualization, G.F.P., T.M. and A.P.; Methodology, M.V., T.C., T.M. and A.P.; Investigation, G.F.P., M.J.S., K.Č., L.A., P.H., M.K., S.K., J.S., M.K., P.P., T.M. and A.P.; Resources, E.F.V., M.Z., H.H., T.H.; Writing – Original draft, G.F.P., M.J.S. and A.P.; Writing – Review & Editing, G.F.P., M.J.S., K.Č., E.F.V., J.H., O.K., P.P., T.M. and A.P.; Visualization, G.F.P., M.J.S., and A.P.; Supervision, O.K., J.H., T.M. and A.P.; Project Administration, T.M., P.P. and A.P.; Funding acquisition, M.J.S., H.H., P.P., T.M. and A.P.

## Data availability

All data described, analysed, and represented in the figures present in this study are available from the corresponding authors upon reasonable request. The mass spectrometry proteomics data have been deposited to the ProteomeXchange Consortium via the PRIDE^73^ partner repository and will be publicly available as of the publication date.

## EXPERIMENTAL MODEL AND STUDY PARTICIPANT DETAIL

### Human cell lines and primary fibroblasts

COX6B1 (6BKO), SDHA (SDHAKO), and ATP5F1B (BETAKO) HEK293 (ATCC CRL-1573) knock-out cells were previously generated^8,74^. For alternative oxidase expression in 6BKO cells (to produce 6BAOX), pcDNA3.1+ mammalian expression vector (Thermo Fisher Scientific, USA) containing the coding sequence of full-length AOX (from *Aspergillus nidulans)*, followed by a C-terminal HA-tag, was transfected as described in^8^. Stably transfected cells were selected with 2 mg/mL G418. The HEK-derived cell lines were maintained at 37 °C and 5% CO_2_ atmosphere in DMEM/F-12 medium (Biowest, L0092) supplemented with 10% (v/v) FBS (Thermo Fisher Scientific, 10270-106), 40 mM HEPES, antibiotics (100 U/mL penicillin + 100 μg/mL streptomycin, Thermo Fisher Scientific, 15140-122) and 50 μM uridine. To assess the impact of hypoxia, the WT cells were cultured in normoxia (21% O2) or placed in a hypoxic chamber (1% O_2_, Xvivo System X3) for 1 week. Label-free quantification mass spectrometry samples, lipid droplets quantification and metabolomic and lipidomic analyses were prepared as described below in a hypoxic chamber.

Fibroblasts from the patients and controls were immortalised as described previously^36^. The cells were cultured in Dulbecco’s modified Eagle’s medium (VWR, 392-0415) supplemented with 10% (v/v) FBS (Thermo Fisher Scientific, 10270-106), 20 mM HEPES and antibiotics (100 U/mL penicillin + 100 μg/mL streptomycin, Thermo Fisher Scientific, 15140-122).

### Human plasma

Following an overnight fast, blood samples were drawn from the antecubital vein of patients and healthy controls between 7:00 and 8:00 a.m. Plasma was collected by centrifugation (2000 RPM RT, 10 min) from one mL EDTA-treated blood and aliquots were stored at −80 °C until lipidomic analyses. A total of 13 patients with MERRF syndrome and 23 age-matched healthy controls were included. The study was performed in accordance with The Code of Ethics of the World Medical Association (Declaration of Helsinki), and the study protocol was approved by the Ethical Review Board of the First Faculty of Medicine and General University Hospital in Prague (77/21 Grant AZV VES 2022 VFN, 17.6.2021). Written informed consent was obtained from all subjects.

## METHODS

### Metabolomic, lipidomic and fluxomic analysis

#### Sample preparation and extraction

HEK-derived cells were seeded in triplicate on 6-well plates (600t/well) and cultured in the growth media (see above) for 24h. For profiling, the cells were rinsed twice in ice-cold PBS, the medium was aspirated, and the cells were immediately frozen at −80°C. For fluxomics, the cultured medium was replaced with DMEM/F-12 medium (Biowest, L0091) supplemented with 10% (v/v) dialysed FBS (Thermo Fisher Scientific, 26400044), 2 mM L-glutamine (Thermo Fisher Scientific, 25030-029), 10 mM glucose (Thermo Fisher Scientific, A24940-01), 40 mM HEPES, antibiotics (100 U/mL penicillin + 100 μg/mL streptomycin, Thermo Fisher Scientific, 15140-122) and 50 μM uridine. After 2 hours, the medium was changed for tracing medium – DMEM/F-12 medium (Biowest, L0091) supplemented with 10% (v/v) dialysed FBS (Thermo Fisher Scientific, 26400044), 40 mM HEPES, antibiotics (100 U/mL penicillin + 100 μg/mL streptomycin, Thermo Fisher Scientific, 15140-122), and 50 μM uridine. The glucose tracing medium was further supplemented with 2 mM L-glutamine and 10 mM ^13^C_6_–glucose (CIL, CLM-3612-1). The glutamine tracing medium with 2 mM ^13^C_5_–glutamine (CIL, CLM-1822-H-0.1) and 10 mM glucose.

Cells were processed using the LIMeX workflow, and polar metabolites were extracted with a biphasic solvent system composed of cold methanol, methyl tert-butyl ether, and 10% methanol, followed by liquid chromatography-mass spectrometry (LC–MS) analysis^75^. Specifically, an aliquot of the upper organic phase was evaporated, resuspended in methanol with the internal standard [12-[(cyclohexylamino) carbonyl]amino]-dodecanoic acid (CUDA), and analysed using lipidomics platforms in positive and negative ion modes. Another aliquot of the upper organic phase was hydrolysed using potassium hydroxide in methanol. The solution was neutralised with hydrochloric acid, and the fatty acids were isolated using hexane. After evaporation, the dry extracts were resuspended in methanol with CUDA and analysed using the lipidomics platform in negative ion mode. Next, one aliquot from the bottom aqueous phase was evaporated and resuspended in an acetonitrile/water (4:1, v/v) mixture containing CUDA and Val-Tyr-Val as internal standards. This sample was analysed using the hydrophilic interaction chromatography (HILIC) metabolomics platform in positive electrospray ionisation mode. Another aliquot of the bottom phase was mixed with an acetonitrile/isopropanol (1:1, v/v) mixture, evaporated, then resuspended in 5% methanol with 0.2% formic acid, again including CUDA and Val-Tyr-Val as internal standards. This sample was analysed using the reversed-phase liquid chromatography (RPLC) platform in negative electrospray ionisation mode.

#### LC – MS analysis and data processing

Six different LC-MS platforms (LIMeX-6D) in positive and negative ionisation modes were used^76^. The LC–MS analysis was conducted using a Vanquish UHPLC system (Thermo Fisher Scientific) coupled to an Orbitrap Exploris 480 mass spectrometer (Thermo Fisher Scientific). Detailed chromatographic and detection parameters are provided in^77^. The data from metabolite and lipid profiling were analysed in Metaboanalyst 5.0 as described in^78^. For ^13^C-metabolic flux analysis, MS1 data were acquired at a mass resolving power of 180,000 FWHM. Data were processed using the MRMPROBS software to provide peak heights of ^13^C-isotopologues based on the total number of carbon atoms per analyte. Isotopic labelling was corrected for natural ^13^C abundance using IsoCor software^79^.

### Proliferation

Studied cell lines were seeded on a 96-well plate (5 x 10^3 cells per well). Where indicated, the cells were pretreated with the mix of 1 μM DGAT1 (A922500, Merck) and 10 μM DGAT2 (PF-06424439, Merck) inhibitors (DGAT1/2i), 200 nM ferroptosis inhibitor ferrostatin-1 (Merck) or 25 μM BSA-conjugated palmitic or oleic acid. After 24 hours, the culture medium was replaced with medium containing pretreatment and either vehicle or 1 μM RSL3 (Merck, SML2234). After 24 and 48 hours, the plate was dumped, blotted and stored at −80°C. Cell proliferation was measured using the CyQUANT Cell Proliferation Assay (Thermo Fisher Scientific, C7026), following the manufacturer’s instructions. Fluorescence intensity at 480/520 nm (ex/em) was recorded using an Infinite M200 plate reader (Tecan Group Ltd., Switzerland), and growth rates were calculated as the fold change in intensity relative to the vehicle. Doubling time (DT) was calculated as: DT = (t2-t1)*log2/log(I_t2_/I_t1_), where I_t1_ and I_t2_ are fluorescence intensities at time t1 and t2 that are proportional to cell number. The data were normalised to WT values. The measurement was performed in six independent experiments.

### Fatty acid oxidation

Cells were seeded on a 24-well plate (200,000/well). After 24h, the cultured medium was replaced with medium containing 100 μM palmitic acid (Merck, P9767), 1 mM carnitine (Merck, C0283) and 1.7 μCi [9,10(n)-^3^H] palmitic acid (GE Healthcare, UK) in the presence or absence of 100 μM etomoxir (Merck, 236020) and the cells were incubated for 4 h. Medium was collected to analyse the amount of released ^3^H_2_O that was formed during the cellular oxidation of [^3^H]-palmitate as described in^80^. The etomoxir-sensitive portion was presented in nCi/mg protein/h. The measurement was performed in 3 independent experiments.

### Fatty acid conjugation

Both sodium palmitate (Merck, P9767) and oleate (Merck, O7501) were conjugated to BSA at 1:6 ratio to the final concentration of 1 mM^81^. Briefly, fatty acids were dissolved in 150 mM NaCl at 70 °C, until no longer cloudy. Then, they were sequentially added to 34 mM BSA in 150 mM NaCl and stirred for 1 hour at 37 °C. Finally, the volume was adjusted with 150 mM NaCl to yield 1 mM final solution, and the pH was adjusted to 7.4.

### Metabolic modelling

To conduct metabolic flux analysis, the INCA software^82^ was used. A simple model, encompassing glycolysis, Krebs cycle, biosynthesis of palmitic acid from acetyl-CoA, gradual biosynthesis of stearic, oleic and palmitoleic acid and, eventually, biosynthesis of triglycerides (TG 50:1, TG 48:1) and phospholipid (POPC) from building blocks defined in previous steps, was defined as an approximation of relevant lipidomic pathways. MS data describing the isotopic enrichment of the fatty acids and lipid molecules mentioned above were introduced to said model, and metabolic fluxes were estimated in INCA to minimise the differences between observed and estimated isotopic enrichment values. To estimate the data published, a ratio of flux of newly synthesised palmitic acid to POPC and to all the lipids considered (POPC and triglycerides) was calculated.

### Lipid droplet imaging

Cells (4– 8 × 10^5^) were seeded on glass coverslips in 6-well plates and incubated for 24 hours. The cells were fixed in 3.7% PFA (Merck, F1635) in warm PBS for 20 minutes, washed twice with PBS and stored at 4°C for a maximum of 2 weeks before imaging. Samples were stained with BODIPY™ 558/568 (1:2000, Thermo Fisher Scientific, D3835) and DAPI (5 μg/mL, Thermo Fisher Scientific, D1306) for 30 minutes, protected from light, and then rinsed twice in PBS for 10 minutes. Z-stack images were collected using a Leica SP8 WLL MP laser scanning confocal microscope (HC PL APO 63x water immersion objective (1.20 N.A.)). The z-step was set to 300 nm, and the confocal zoom was adjusted to maintain a pixel size of approximately 100nm. Images were analysed using the FIJI/ImageJ software^83^. Briefly, z-projected images of each stack were generated, the background was subtracted and thresholded to delimit the signal corresponding to lipid droplets. The “analyse particles” tool was used on the binary thresholded images to quantify the total stained area. The number of nuclei in the DAPI-stained images was manually counted.

### Label-free quantification mass spectrometry analysis

Label-free quantification mass spectrometry analysis (LFQ-MS) of cell pellets was performed in triplicate by the Proteomics Service Laboratory at the Institute of Physiology and the Institute of Molecular Genetics of the Czech Academy of Sciences. Cellular pellets (100 μg of protein) solubilised by sodium deoxycholate (final concentration 1%) were processed according to the ethylacetate extraction in-solution trypsin digestion protocol as described in^84^. About 500 ng of desalted peptide digests were separated on 50 cm C18 column (EASY-Spray ES903, Thermo Fisher Scientific) using 120 min elution gradient (Dionex Ultimate 3000, flowrate 300 nL/min) and acquired in a DIA mode on an Orbitrap Exploris 480 mass spectrometer (Thermo Fisher Scientific). Thermo raw files were processed by the directDIA mode and visualized in Spectronaut 19.9 (Biognosys) software using the default settings with Precursor and Protein Q-value and PEP cutoff set at 0.01. MMTS alkylated cysteine was selected as a fixed modification Methylthio (C). Search was performed against the human reference proteome UP000005640_9606.fasta (UniProt release 2024_01). Protein groups quantities (PG.Quantity, MS2 level) from the Protein report generated by the Spectronaut were evaluated in Perseus v. 2.1.3.0^85^ and further visualised in GraphPad Prism.

### Enzyme activities

Complex IV and citrate synthase (CS) activities were determined spectrophotometrically^86^ at 30 °C in cell lysates. The complex IV assay medium consisted of 40 mM K-Pi, 1 mg/ml BSA, and a pH of 7.0. The reaction was started with 30 μM reduced cytochrome *c*, and its oxidation was monitored at 550 nm for 40 s. Citrate synthase (CS) activity was determined using a medium containing 0.1 M Tris–HCl, 0.1 mM 5,5′-dithiobis-(2-nitrobenzoic acid), 50 μM acetyl coenzyme A, pH 8.1. The reaction was initiated by adding 0.5 mM oxaloacetate and then monitored for changes at 412 nm for 1 min. The data were corrected for the absorbance change in the absence of oxaloacetate. Enzyme activities were expressed as nmol/min/mg protein using molar absorption coefficient ε_550_ = 19.6 mM/cm (complex IV) or ε_412_=13.6 mM/cm (CS).

### Evaluation of metabolic fluxes

Seahorse Extracellular Flux (XF) Analyser (Agilent Technologies, USA) was used to assess mitochondrial respiration and glycolytic rate in parallel, as described before^8^. Briefly, 2-3.5 × 10^4 cells were seeded on poly-L-lysine-coated 24-well measuring plate and incubated overnight under standard conditions. Before the evaluation, the cells were rinsed with 1mL of measuring media (DMEM (Merck, D5030; pH 7.4, 37°C) supplemented with 0.2% (w/w) BSA and 1 mM pyruvate). Cells were incubated in measuring medium for 30 minutes at 37°C. The oxygen consumption rate (OCR) and extracellular acidification rate (ECAR) were recorded at basal metabolic rate and after subsequent additions of 10 mM glucose, 1μM oligomycin, 1μM FCCP, and a mixture of 1μM rotenone, 1 μg/mL antimycin A, 100 mM 2-deoxyglucose (100 mM) and 5 μg/ml Hoechst 33342. The OCR and ECAR were normalised to the cell count. The number of cells in each well was counted using Cytation 3 Cell Imaging Reader (BioTek) and analysed using the Gen5 software (BioTek). Sample evaluations were performed in at least three independent experiments.

### NAD^+^/NADH

Cells (10 × 10^3^ per well) were seeded in 96-well plates, and the ratio of nicotinamide adenine dinucleotide (NAD^+^/NADH) was assessed using the NAD^+^/NADH Glo assay (Promega, Madison, WI, USA), following the manufacturer’s protocol for seeded cells.

### GSH detection assay

Cells (5 × 10^3^ per well) were seeded in 96-well plates, and the GSH level was assessed using the GSH/GSSG Glo assay (Promega, Madison, WI, USA), following the manufacturer’s protocol for seeded cells.

### Data analysis, visualisation, and statistics

Figures were prepared and statistics were calculated using GraphPad PRISM 10. Data are presented as means with error bars representing the standard errors, unless otherwise indicated. One-sample t-test, one-way ANOVA and 2-way ANOVA tests were utilised to calculate p values; a p value of less than 0.05 was considered significant. Statistical details can also be found in the figure legends.

