## Supplemental figures for "Sequestration of the polyunsaturated fatty acids protects the cells with oxidative phosphorylation deficiency from ferroptosis"

[1]*Laboratory of Bioenergetics, Institute of Physiology, Czech Academy of Sciences, Vídeňská 1083, Prague, Czech Republic*
[2]*Faculty of Science, Charles University, 12800 Prague, Czech Republic*
[3]*Laboratory of Metabolism of Bioactive Lipids, Institute of Physiology, Czech Academy of Sciences, Vídeňská 1083, Prague, Czech Republic*
[4]*Laboratory of Metabolomics, Institute of Physiology, Czech Academy of Sciences, Vídeňská 1083, Prague, Czech Republic*
[5]*Department of Biochemistry and Molecular and Cellular Biology. Faculty of Health and Sport Sciences. University of Zaragoza, Spain*
[6]*IRCCS Women and Children's Hospital "Burlo-Garofolo", Trieste, Italy*
[7] *Department of Paediatrics and Inherited Metabolic Disorders, First Faculty of Medicine, Charles University, General University Hospital, Prague, Czech Republic.*
[*] Contributed equally
[#] corresponding author

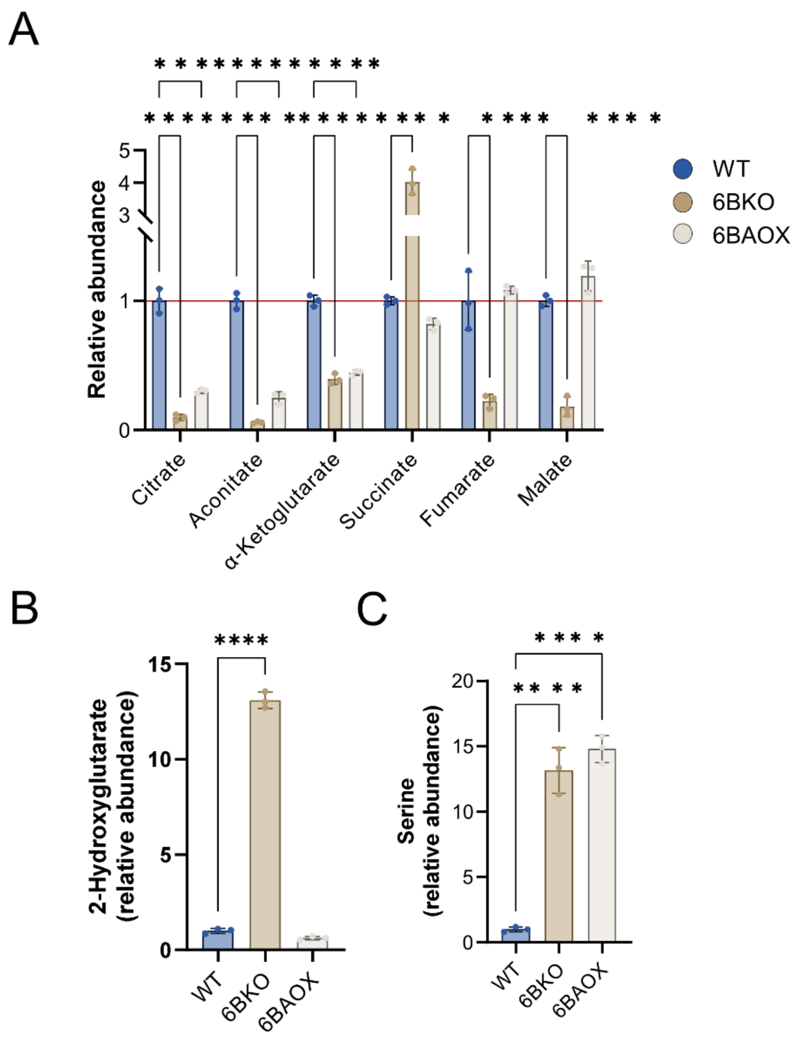


**Figure S1: Metabolic remodelling in CIV-deficient cells**

Metabolic profiling (LC-MS) of intracellular TCA cycle metabolites (A), 2-hydroxyglutarate (B) and serine (B) after 24h incubation in fresh culture media. Data are expressed as fold changes compared to WT (n = 3). Bar graphs represent means ± SD. One-way (B, C) or 2-way (A) ANOVA was performed (*p*-value: * <0.05; ** <0.0.1; *** <0.001; **** <0.0001).

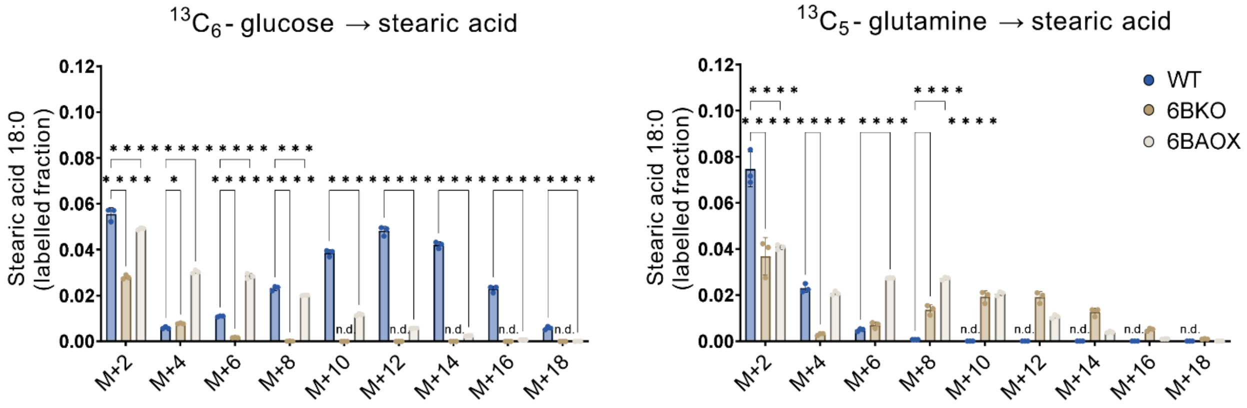


**Figure S2: 13C labelling of fatty acids**

$^{13}$C labelling of stearic acid after 24h incubation with $^{13}C_6$-glucose or $^{13}C_5$-glutamine measured by lipidomic analysis after fatty acyl chain lipolysis (n = 3). Bar graphs represent means ± SD. 2-way ANOVA was performed (*p*-value: * <0.05; ** <0.0.1; *** <0.001; **** <0.0001).

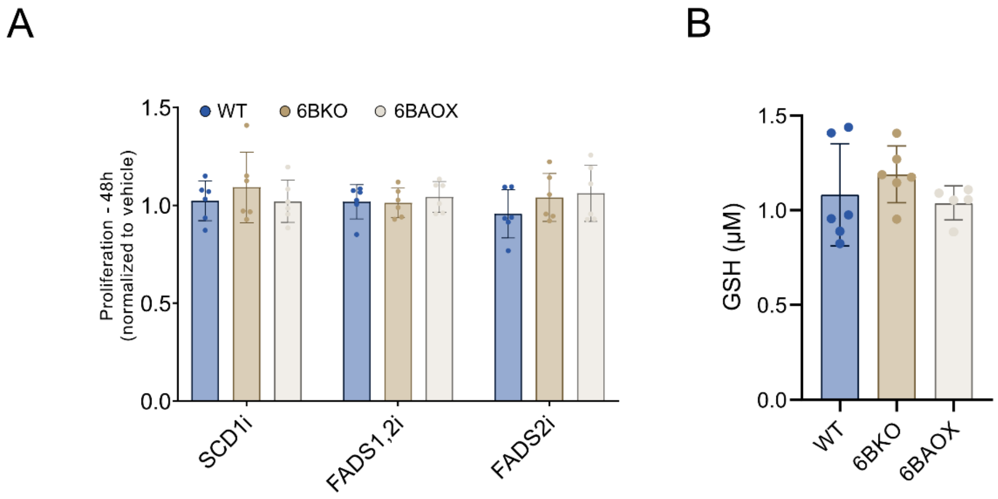


**Figure S3: Proliferation and GSH level in CIV-deficient cells**

(A) Cell proliferation analysis of the cell lines incubated for 48h with inhibitors against SCD1 (SCD1i), FADS1 and 2 (FADS1,2i). Data are expressed as fold change to vehicle (n = 6). (B) (E) GSH level assessed in WT, 6BKO (n =6). Bar graphs represent means ± SD. 2-way ANOVA was performed (*p*-value: * <0.05; ** <0.0.1; *** <0.001; **** <0.0001).

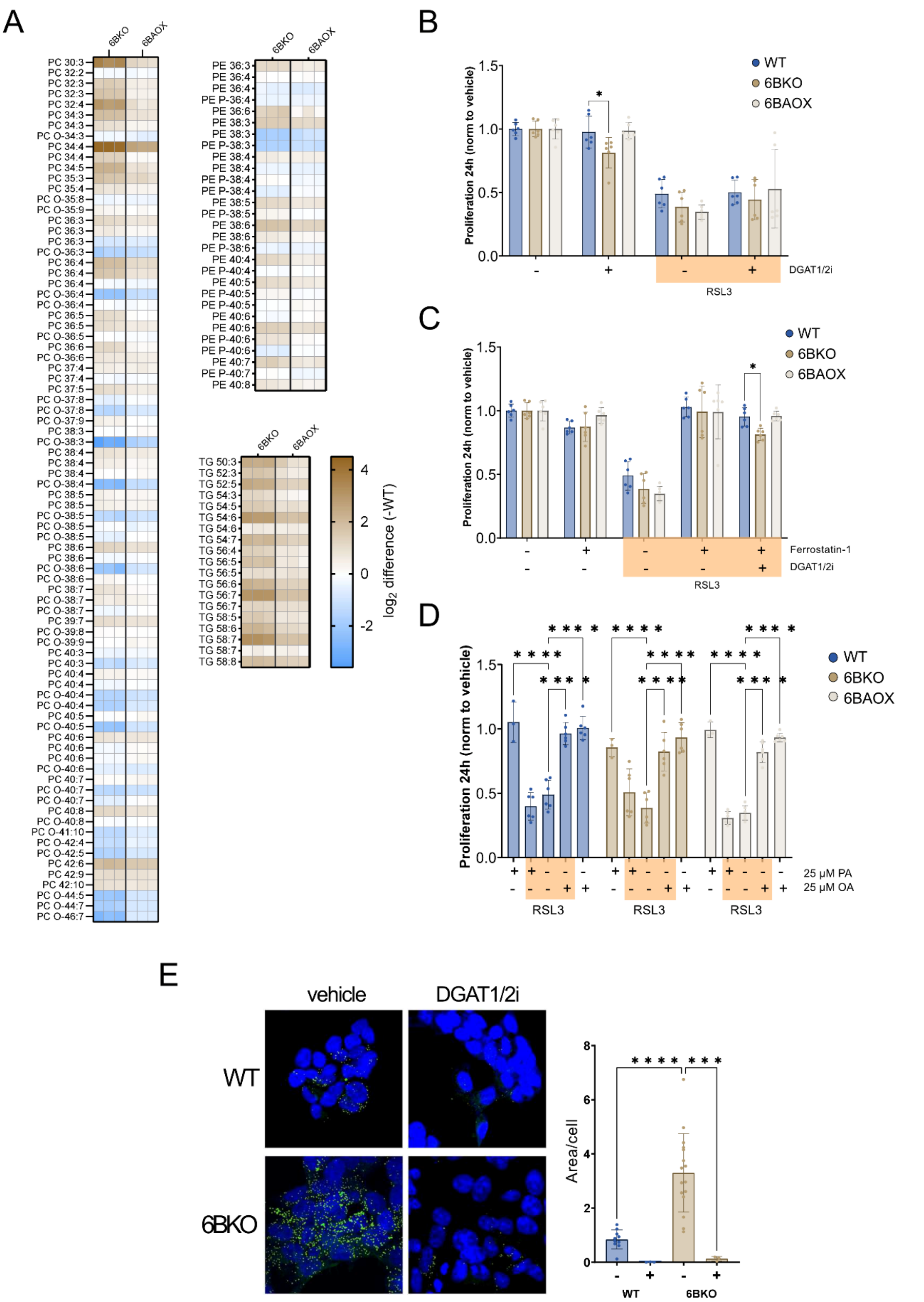


**Figure S4: PUFA-mediated stress response in CIV-deficient cells**

(A) The heatmaps showing relative levels of PC, PE and TG containing at least one PUFA in 6BKO and 6BAOX. Data are expressed as $log_2$(fold change) compared to WT (n = 3). (B) Cell proliferation analysis of the cell lines incubated for 24h with inhibitors against GPX4 (RSL3 1µM). The cells were pretreated for 24 hours with a mixture of DGAT1 (A922500, 1 µM) and DGAT2 (PF-06424439, 10 µM) inhibitors

(DGAT1/2i) (B, C), ferroptosis inhibitor ferrostatin-1 (200 nM) (C), or BSA-conjugated palmitic (PA) or oleic (OA) acid (D). Data are expressed as mean ± SD of fold change to vehicle (n = 6). (E) Representative image and quantification of BODIPY 493/503 staining of the WT, 6BKO cell lines (n ≥ 3) treated with vehicle or DGAT1 and DGAT2 inhibitors. Bar graphs represent means ± SD. One-way (E) or 2-way (B-D) ANOVA was performed (*p*-value: * <0.05; ** <0.0.1; *** <0.001; **** <0.0001).

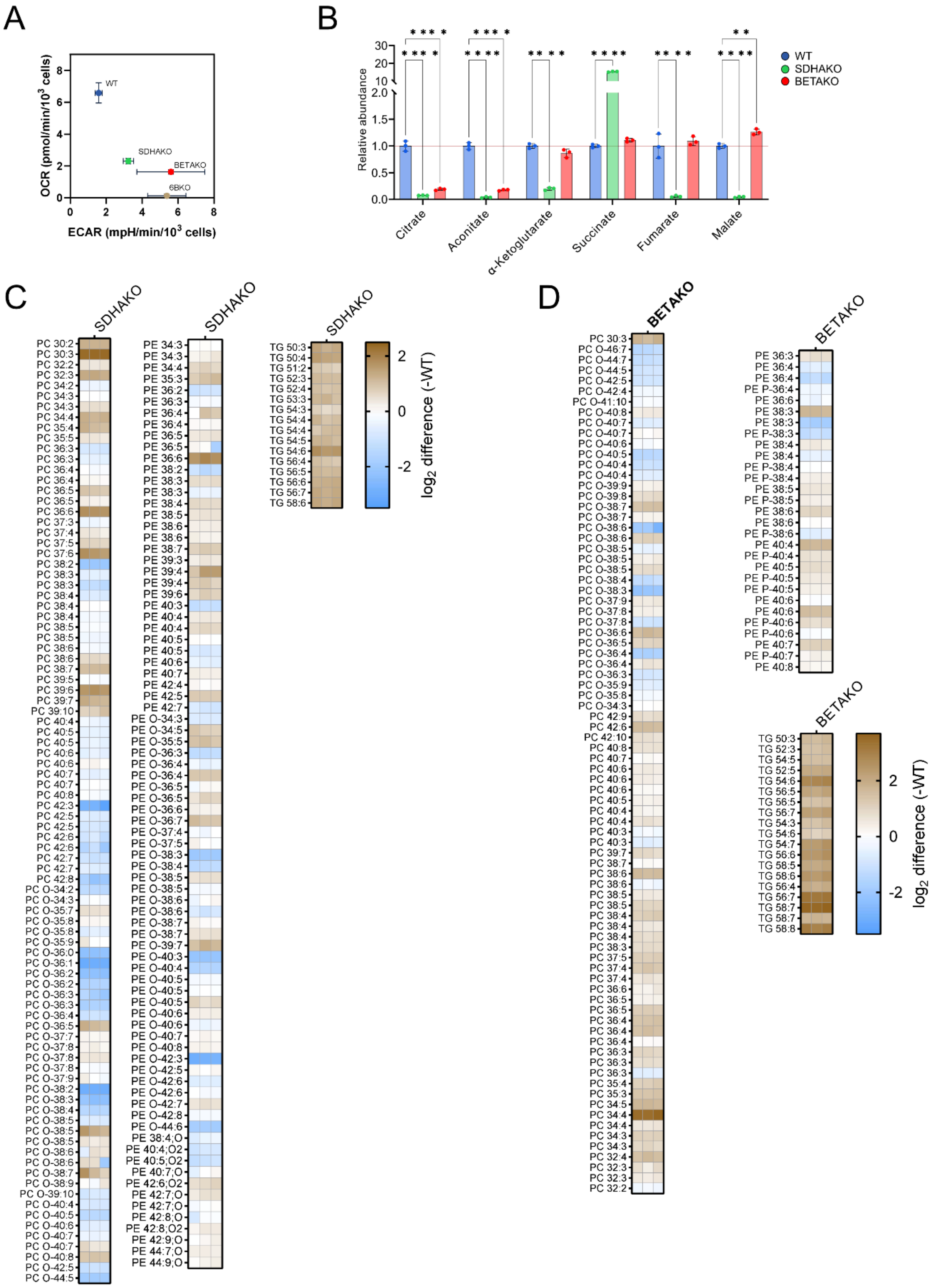


**Figure S5: Metabolic remodelling in CII- and CV-deficient cells**

(A) Metabolic phenotype of WT, 6BKO, BETAKO (data published in Cunatova24) and SDHAKO represented as a phenogram – combined parallel evaluation of cellular oxygen consumption rate

(OCR) and extracellular acidification rate (ECAR) (mean ± SEM, n ≥ 3). (B) Metabolic profiling (LC-MS) of TCA cycle metabolites after 24h incubation in fresh culture media. Data are expressed as fold changes compared to WT (means ± SD, n=3). Asterisks represent *p*-value: * <0.05; ** <0.0.1; *** <0.001; **** <0.0001. (C, D) The heatmaps showing relative PC, PE and TG levels containing at least one PUFA in SDHAKO (C) and BETAKO (D). Data are expressed as $log_2$(fold change) compared to WT (n = 3). Bar graphs represent means ± SD. 2-way ANOVA was performed (*p*-value: * <0.05; ** <0.0.1; *** <0.001; **** <0.0001).

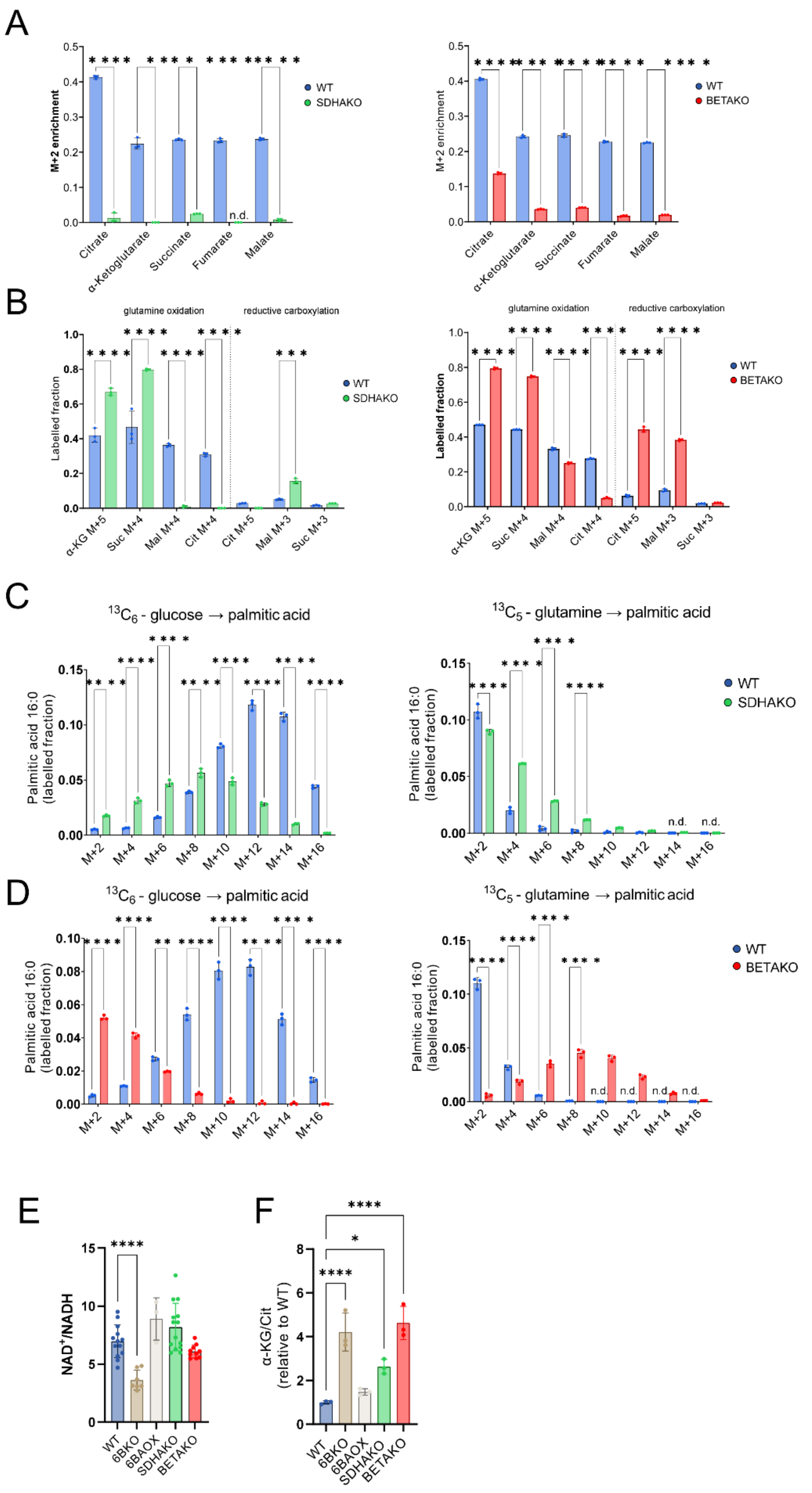


**Figure S6: Metabolic remodelling in CII- and CV-deficient cells**

The WT, SDHAKO and BETAKO cells were incubated with the tracer ($^{13}C_6$-glucose or $^{13}C_5$-glutamine) for 24h, and the labelled fraction of M+2 TCA cycle metabolites derived from $^{13}C_6$-glucose (A) or $^{13}C_5$-glutamine (B) was assessed by LC-MS (n = 3). $^{13}C$ labelling of palmitic acid from $^{13}C_6$-glucose or $^{13}C_5$-

glutamine in SDHAKO (C) or BETAKO (D) cells measured by lipidomic analysis after fatty acyl chain lipolysis (n = 3). (E) $NAD^+$/NADH ratio assessed in WT, 6BKO, BETAKO (data published in[8]) and SDHAKO (n ≥ 3). (F) α-ketoglutarate/citrate ratio measured by LC-MS after 24h incubation in fresh culture media. Data are expressed as fold changes compared to WT (n = 3). Bar graphs represent means ± SD. One-way (E, F) or 2-way (A – D) ANOVA was performed (*p*-value: * <0.05; ** <0.0.1; *** <0.001; **** <0.0001).
